# Mapping the complete influenza A virus infection cycle through single vRNP imaging

**DOI:** 10.1101/2025.01.20.633851

**Authors:** Huib H. Rabouw, Janin Schokolowski, Micha Müller, Matthijs J.D. Baars, Antonella F.M. Dost, Theo M. Bestebroer, Jakob Püschel, Hans Clevers, Ron A.M. Fouchier, Marvin E. Tanenbaum

## Abstract

Cell-to-cell heterogeneity is a common feature of viral infection that can generate enormous complexity, complicating understanding of infection progression and interpretation of differences between viral variants. To overcome these challenges, we developed a technology to visualize the infection cycle of human influenza A virus (IAV) from start-to-finish in individual living cells with unprecedented spatial and temporal resolution, which identified numerous distinct pathways through which infection can progress. We show that heterogeneous viral gene expression drives infection cycle heterogeneity, and identified genome packaging, vRNP transcriptional activation and host cell division as major determinants of gene expression variability. Our work maps out the complete IAV infection cycle, identifies the origin of infection heterogeneity and provides a broadly-applicable technology for studying IAV and other viruses.

## Introduction

Viral infections typically proceed through a well-ordered series of molecular events, which include host cell entry, intracellular trafficking, viral transcriptional activation, viral protein synthesis, genome replication, followed by genome packaging and budding of new virions that go on to infect other cells or individuals. While the canonical infection cycle of a number of model viruses has been relatively well-described, viral infection can be highly heterogeneous at a single-cell level in terms of kinetics, success, and host cell responses [1-4]. For example, many studies have shown that only a small subset of virions in a viral inoculum typically cause (detectable) infection in cells in culture [5-7], but the alternative infection cycle pathways, as well as the mechanisms of infection heterogeneity are mostly unknown. The lack of knowledge on infection cycle heterogeneity is largely due to limitations of current methodology in visualizing infection in individual living cells with sufficient resolution. Here, we develop an approach for single-molecule live-cell imaging of infection and apply this approach to generate a comprehensive and quantitative single-cell description of the entire infection cycle of Influenza A virus (IAV), a highly-studied negative-strand RNA virus that is an important human and animal pathogen. Our analysis provides the first kinetic map of infection, uncovers numerous different ‘off-pathway’ infection cycles through which infection can progress, identifies key bottlenecks that cause abortive infection and reveals the origin of heterogeneity in the host cell antiviral response.

## Results and Discussion

### Visualizing early IAV infection by single-molecule imaging

To map IAV infection cycle dynamics (Figure 1A), we aimed to develop a technology that would allow visualization of IAV infection in living cells with single vRNP resolution. The IAV genome is made up of eight negative-strand viral RNAs (vRNA), called genome segments. Each vRNA is encapsidated by dozens of copies of the viral nucleoprotein (NP) protein and is bound by the trimeric viral polymerase complex, consisting of the PA, PB1 and PB2 subunits, together called the viral ribonucleoprotein (vRNP). To label single vRNPs in living cells, we expressed a fluorescently-tagged anti-NP nanobody fused to a nuclear localization signal (Nb^NP^-GFP) in human A549 cells and infected cells with wildtype IAV (strain PR8) (Figure 1B) [8]. Bright fluorescent foci were observed in individual cells by spinning-disk confocal microscopy after virus inoculation, but not in mock inoculated cells (Figure 1C, S1A and S1B), nor in cells pre-treated with the IAV entry inhibitor bafilomycin A1 (BafA1) (Figure S1C) [9]. At low multiplicity of infection (MOI), infected cells typically contained 7 or 8 spots, consistent with the expected number of vRNPs per virion (Figure S1D), while >8 foci were observed at higher MOI, allowing classification of cells as uninfected, or infected with 1, 2, or >2 virions (Figure 1C and 1D). Since this approach enabled single vRNP imaging of negative-strand RNA viruses, we refer to this method as Virus Infection Real-time Imaging Negative-strand (VIRIM-neg), in line with the ‘VIRIM’ terminology, a method to visualize single positive-strand RNA viruses [10]. Combining VIRIM-neg with single-molecule fluorescence *in situ* hybridization (smFISH) labeling of vRNAs revealed that the large majority of VIRIM-neg foci represented vRNPs, with a small subset of VIRIM-neg spots likely representing encapsidated copy RNAs (cRNAs), positive-strand genome replication intermediates (Figure S1E and S1G). Importantly, labeling of vRNPs by VIRIM-neg did not detectably alter virus fitness (Figure S1H and S1I). VIRIM-neg spots represent full-length genome segments, rather than truncated defective interfering genomes (DIGs) [11] (Figure S1J). Importantly, VIRIM-neg was successfully applied to a range of phylogenetically distant IAV isolates from human, swine, and avian origins, and was even successful for patient-derived virions (Figure 1E), enabling comparison of the infection cycle of different IAV strains and isolates.

**Figure 1.**
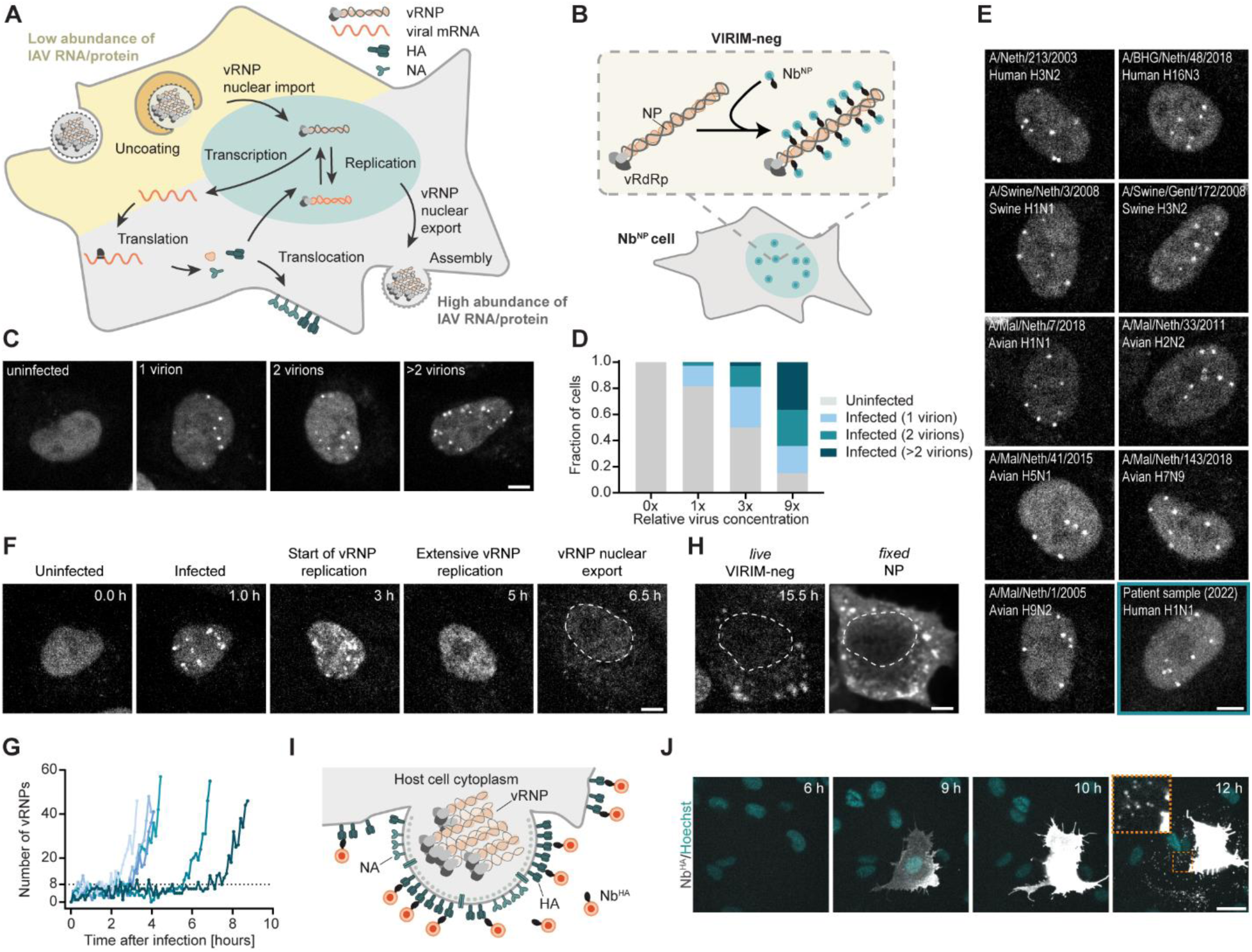
New technologies to image single IAV vRNPs and IAV protein in live cells. **(A)** Cartoon of the IAV infection cycle. Early (yellow shading) and late (grey shading) phases of infection are characterized by low and high abundance, respectively, of viral material in the cell. **(B)** Cartoon of VIRIM-neg single vRNP imaging assay. Upon entry, fluorescently labeled Nb^NP^ molecules bind to multiple copies of NP present in vRNPs, locally enriching fluorescence signal. Nb^NP^ fluorescent spots in cells thus represent single vRNPs. **(C)** Example images of A549-Nb^NP^ cells inoculated with IAV (strain PR8) at 1 hpi. Shown are examples of cells infected by 0, 1, 2, or >2 virions, inferred from the number of vRNPs. **(D)** Quantification of the frequency of infections with 0, 1, 2, or >2 virions in A549-Nb^NP^ cells 1 hpi at different virus concentrations. **(E)** Example images of A549-Nb^NP^ cells inoculated with the indicated IAV isolates, imaged at 1 hpi. **(F)** Example images from a time-lapse movie of A549-Nb^NP^ cells inoculated with IAV. Images show indicated infection cycle phases, as visualized by VIRIM-neg. **(G)** Quantification of VIRIM-neg spot count over time in infected A549-Nb^NP^ cells. Every line represents a single cell. All traces were aligned in silico to the moment of first detection of VIRIM-neg spots. The expected value of eight vRNPs for a single-virion infection is indicated by the dashed line. Five representative cells are shown in which viral genome replication occurs. **(H)** Last timepoint of the time-lapse movie shown in (G) showing nuclear export of vRNPs by VIRIM-neg (left) and NP immunofluorescence staining (right) of the same cell. **(I)** Cartoon illustrating the extracellular HA and progeny labeling method using the Nb^HA^. Red circles indicate a fluorescent protein associated with the Nb^HA^. **(J)** Example images from a time-lapse movie of an IAV infected cell labelled with Nb^HA^. Orange dashed square indicates zoom-in that highlights viral progeny release. Scale bar, 25 µm. See also Video S2. **(C, E, F, H)** Scale bars, 5 µm. See also Video S1 and S2.

After vRNP entry, the number of VIRIM-neg spots in the nucleus typically remained constant for several hours, after which an increase in numbers was observed, indicative of vRNP replication (Figure 1F, 1G, and S2A). Spot number typically rapidly increased until too many spots were present to resolve spots individually, resulting in a diffuse nuclear signal (Figure 1F, referred to as "Extensive vRNP replication”). The increase in the number of VIRIM-neg spots was matched by an increase in vRNAs, as detected by smFISH, indicating that it reflects bona fide genome replication (Figure S2, B and C). At late stages of infection, cells harboring replicating virus often showed translocation of VIRIM-neg signal from the nucleus to the cytoplasm (Figure 1F (last time point) and 1H), which reflects vRNP nuclear export (Figure 1H and S2D), which is known to occur late in infection to allow genome packaging into new virions at the plasma membrane. In conclusion, we have established VIRIM-neg as a single-molecule live-cell assay to visualize IAV infection, vRNP replication, and vRNP nuclear export. VIRIM-neg does not require genetic modifications of IAV and can be applied to phylogenetically distant IAV variants. As such, VIRIM-neg provides a unique opportunity to study viral infection in single cells.

### Mapping late-stage viral infection

To map the IAV infection cycle from beginning (vRNP entry) to end (viral progeny production), we aimed to develop a second, orthogonal approach to study infection progression during late stages of infection. We purified a fluorescently-labeled Nb targeting the HA extracellular domain (Nb^HA^) (Figure S3A) [12] and added it to the cell culture medium during long-term VIRIM-neg imaging (Figure 1I). Nb^HA^ labeled infected cells from ∼5 hours post inoculation (hpi) (Figure S3B and S3C), allowing real-time, quantitative analysis of viral protein synthesis over the course of infection in single cells without genetic modification of the virus. Notably, most cells that underwent genome replication, as assessed by VIRIM-neg, showed strong Nb^HA^ labeling, but weak staining was observed in a subset of infected cells in which no genome replication was observed, which likely represents viral transcription in the absence of replication (Figure S3D and S3E). Nb^HA^ also successfully labeled single virions, which are coated by HA protein, both for purified virions, and *in situ* in infected cell cultures, allowing real-time single-single cell visualization of viral progeny production (Figure 1J, and S3E to S3G). In summary, Nb^HA^ live-cell imaging is complimentary to VIRIM-neg, and together, these techniques can be used to map the kinetics and cell-to-cell heterogeneity of the IAV infection cycle from start-to-end.

### Single-cell viral infection kinetics

Having established VIRIM-neg and Nb^HA^ live-cell imaging technologies, we set out to generate a detailed map of IAV infection dynamics. Starting at vRNP entry, we first determined the time from virus inoculation to vRNP entry, which we found to be highly heterogeneous among cells, ranging from approximately 10 min to several hours (average 190 ± 169 min) (Figure 2A). To identify the origin of heterogeneity in vRNP entry dynamics, we examined the timing of individual sub-steps in the viral entry pathway in more detail. IAV virions first undergo endocytosis by host cells, followed by virion fusion with endosomal membranes and nuclear import of vRNPs. To measure the time from endosomal uptake to nuclear entry, we synchronized virus entry by exploiting IAV’s dependency on low pH for fusion (Figure S4) [13]. After pH adjustment, vRNPs rapidly accumulated in the nucleus with a median time of 6.1 min (Figure 2A), much shorter than the median time of 140 min needed from virus inoculation to nuclear entry. Therefore, we conclude that the pre-fusion phase (host cell attachment and endocytosis) is the most time-consuming phase of vRNP entry. To dissect the kinetics of post-fusion vRNP release into the cytoplasm, we performed VIRIM-neg imaging with high temporal resolution after synchronized entry. Shortly after pH adjustment a single bright, largely stationary spot appeared in many cells (median time to spot appearance 2.9 min), which likely represent assemblies of all vRNPs in a bundle that are labeled by cytoplasmic Nb^NP^ immediately after endosomal pore formation (Figure 2B and 2C) [14], indicating that endosomal acidification and membrane fusion occurs within minutes. From these bright foci, individual vRNPs split off in a step-by-step fashion at an increasing rate for each subsequent vRNP (Figure 2B and 2D, Video S3), suggesting that the vRNP bundle is becoming destabilized with each released vRNP. The entire vRNP unpacking process is relatively fast (median time of 0.9 min), indicating that this step is not a kinetic bottleneck in IAV infections (Figure 2E). After initial vRNP disassembly, no further interactions were observed between vRNPs in the cytoplasm, resolving the controversy whether vRNPs travel from the endosome to the nucleus separately or bundled [15, 16]. Based on these results, we calculated the duration of intracellular transport of vRNPs from the endosome release site to the nucleus to be ∼2 minutes. Together, our results provide a detailed map of IAV entry dynamics and identify viral attachment to the host cells and the early stages of endocytosis as the major kinetic bottleneck in the viral entry process. We next aimed to map the kinetics of viral infection progression after vRNP nuclear entry. For this, we performed long-term VIRIM-neg and Nb^HA^ imaging, and determined the time between 1) vRNP entry and initiation of replication, 2) initiation of replication and vRNP nuclear export, 3) initiation of replication and HA membrane accumulation and 4) HA membrane accumulation and virion production. Substantial cell-to-cell heterogeneity was observed for most steps of the infection cycle (Figure 2F to 2J). Notably, the step from vRNP nuclear entry to initiation of replication (i.e. synthesis of sufficient viral polymerase and NP) took almost as long as the time from the first replication event to vRNP nuclear export, which encompasses synthesis of 10,000s-100,000s of vRNAs [17] (3.3 vs 5.2 h) (Figure 2H). These measurements reveal that the preparation for the first replication rounds is relatively time-consuming, but once all components are in place, subsequent rounds of replication can occur very fast, illustrating the highly exponential nature of replication during infection. The time from vRNP nuclear export to budding of progeny virions was surprisingly short (∼0.3 h), indicating that vRNP export from the nucleus is precisely timed during infection such that all components for packaging and virion budding have been prepared at the moment when vRNP nuclear export initiates (calculated from Figure 2H to 2J). Late nuclear export of vRNPs may allow maximal accumulation of vRNPs in the nucleus before packaging occurs to generate high numbers of new progeny. Together, these analyses establish a detailed single-cell kinetic map of infection progression and identify host cell attachment and endocytosis, initiation of replication and vRNP nuclear export as major kinetic bottlenecks in infection.

**Figure 2.**
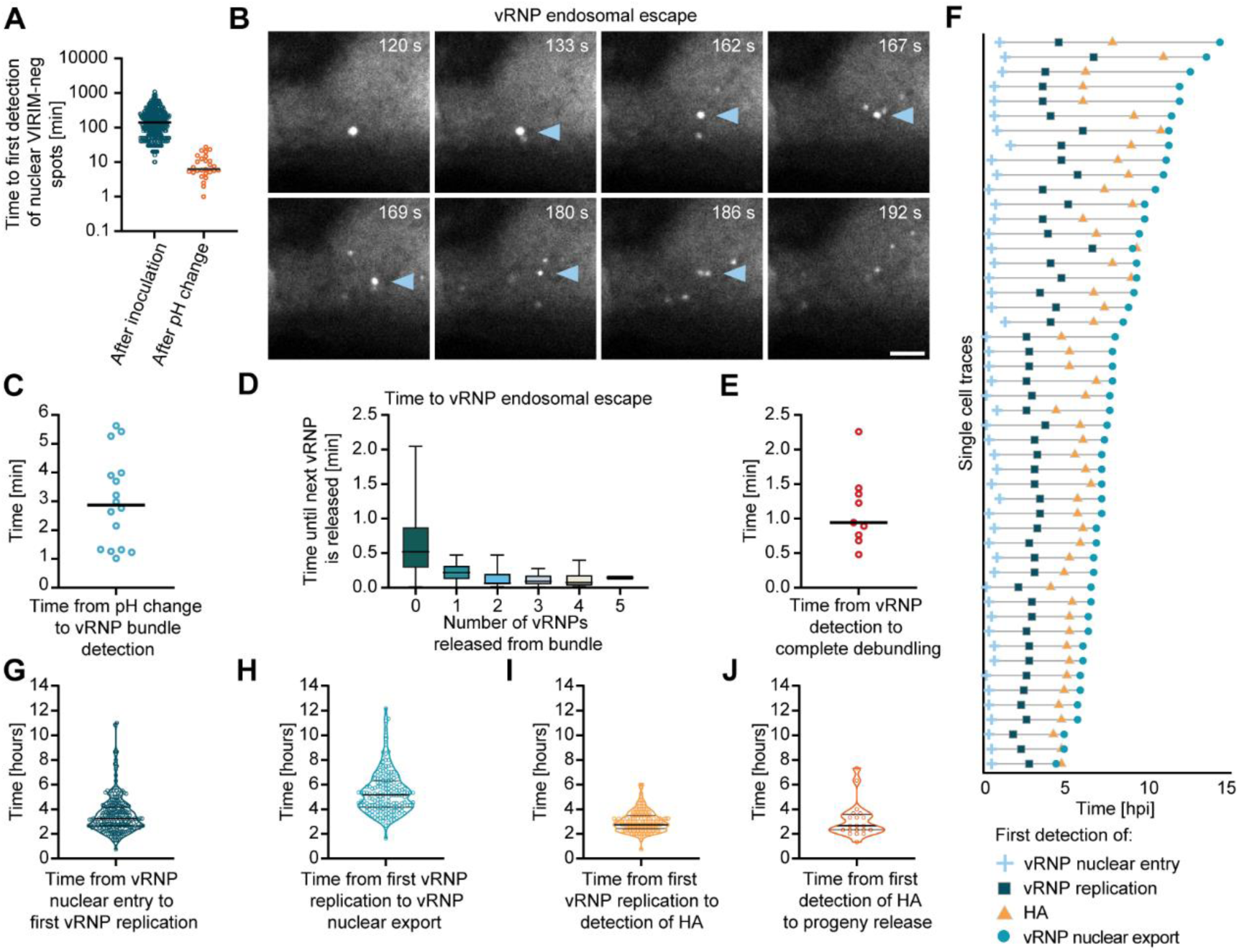
Mapping IAV infection cycle kinetics. **(A)** Quantification of the time of first nuclear detection of VIRIM-neg spots relative to the moment of inoculation (left) or to the moment of pH change (which induces virion-endosome fusion) (right). Dots represent single cells. **(B)** Example images of a time-lapse movie of IAV inoculated A549-Nb^NP^ cells showing vRNP cytoplasmic entry. Arrowheads indicate the slow-moving vRNP assembly, from which single vRNPs split off over time (dim spots). Scale bar, 5 µm. **(C)** Quantification of the time from pH adjustment to first detection of the vRNPs by VIRIM-neg. Dots represent single virions. **(D)** Quantification of the time until the next vRNP is released from the vRNP bundle, relative to the number of released vRNPs. **(E)** Quantification of the time between the first detection of the virion bundle and complete disassembly of the vRNP bundle. Dots represent single virions. **(F)** Example single-cell viral infection cycle traces. The moment of vRNP nuclear entry, initiation of vRNP replication, first HA detection, and vRNP nuclear export are indicated. t=0 indicates the moment of inoculation. **(G)** Quantification of the time between vRNP nuclear entry and vRNP replication initiation. **(H)** Quantification of the time between vRNP replication initiation and vRNP nuclear export. **(I)** Quantification of the time between vRNP replication initiation and HA detection at the plasma membrane. **(J)** Quantification of the time between HA detection at the plasma membrane and progeny virion release. **(G, H, I, J)** Dots represent single cells. See also Video S3.

### Infections progress through diverse pathways

To determine if all infections progress through the same series of molecular events, we next asked whether each step was successfully completed in every infected cell. For this, we compared the viral titer as determined by VIRIM-neg, which assesses the number of virions that enter cells, with the titer as determined by TCID_50_ assay, which assesses the number of virions that cause infections which produce viral progeny (Figure 3A). This comparison revealed a striking ∼100-fold higher viral titer for vRNP entry, suggesting that the vast majority of infecting virions do not go on to produce viral progeny. The surprisingly low success-rate of IAV infection progression was observed irrespective of the method of virus production (Figure 3A), confirming the validity of these results. To determine which step(s) in the infection cycle represent bottlenecks towards successful progeny formation, we first determined the efficiency of vRNP nuclear import, which revealed that vRNPs rarely, if ever, fail to translocate from the cytoplasm to the nucleus (Figure 3B and 2C, Videos S4 and S5). Thus, vRNP nuclear import does not represent a major bottleneck for successful infection. In contrast, using long-term VIRIM-neg and Nb^HA^ imaging, we found that every step after vRNP nuclear import that we assessed (genome replication, viral protein synthesis, vRNP nuclear export and virion production) failed in a considerable fraction of infected cells, allowing us to distinguish six distinct classes of infection progression pathways (Figure 3D and 3E). Thus, the infection cycle of IAV contains numerous bottlenecks, explaining the low overall success-rate of IAV infections. To assess viral infection dynamics and heterogeneity in a physiologically-relevant context, we first repeated VIRIM-neg assays in airway epithelial organoids; primary, untransformed cell cultures that accurately recapitulate cell types found in the human airway (Figure S5A) [18]. We found very similar dynamics and success-rates of replication (Figure S5B and S5C), as well as HA synthesis (Figure S5D and S5E) in airway organoids as compared to A549 cells, confirming the validity of our findings in A549 cells (Figure 3F).

**Figure 3.**
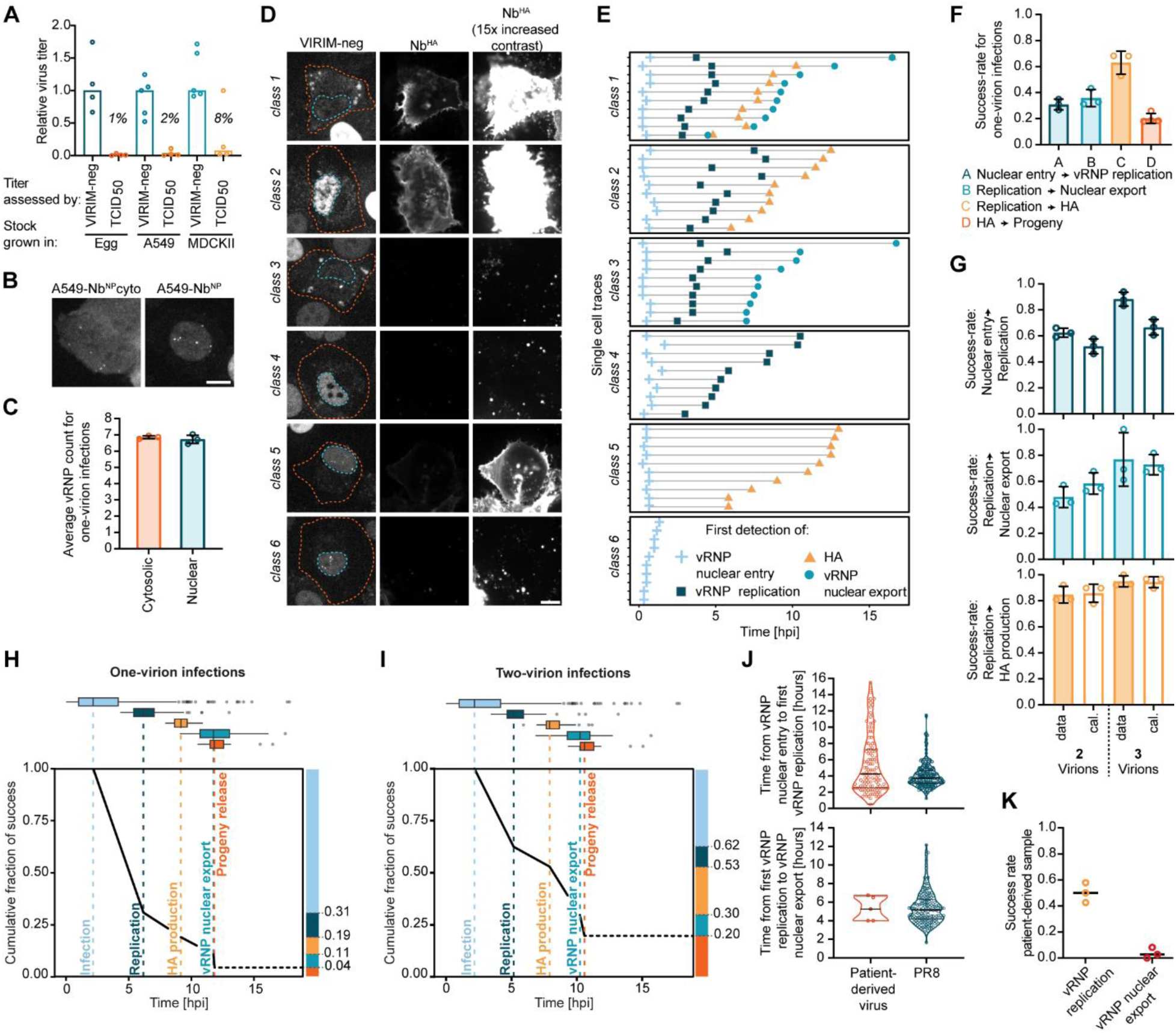
Identification of bottlenecks in IAV infection. **(A)** Viral titer for virus stocks produced in either A549 cells, MDCKII cells or in chicken eggs. The titer was determined either by VIRIM-neg, which detects vRNPs after entering the host cell nucleus, or by TCID_50_, which detects only successful infections that cause detectable cytopathic effect. Titers are shown relative to the median VIRIM-neg titer. Dots represent independent experiments. **(B)** Example images of VIRIM-neg spots either in the cytoplasm of A549-Nb^NP^cyto cells or in the nucleus of A549-Nb^NP^ cells, imaged at 1 hpi. Scale bar, 10 µm. **(C)** Quantification of the number of vRNPs delivered into the cytoplasm or nucleus. **(D)** Example images for six IAV infection phenotype classes. Orange dashed lines indicate cell outlines. Blue dashed lines indicate nuclear outlines. All Nb^HA^ images are also shown with 15-fold increased contrast (right panels). Note, that at these contrast settings, virions and fluorescent cell debris is also visible. Scale bar, 10 µm. **(E)** Example single-cell infection cycle traces, which indicate the moment of nuclear vRNP detection, initiation of vRNP replication, first HA detection, and vRNP nuclear export. Each horizontal line represents a single cell. (F) Quantification of average fraction of one-virion infected cells in which vRNP replication, vRNP nuclear export, HA production, and progeny release are successful. **(G)** Experimental (data) and calculated fraction of cells (cal.) with successful vRNP replication (top), vRNP nuclear export (middle) or HA production (bottom) for infections by 2 or 3 virions. Expected success rates were calculated assuming that each virion provided an independent chance of success (see methods). **(H,I)** Graphs summarizing the kinetics and success rates of each infection cycle phase for one-virion (H) or two-virion (I) infections. Median times are indicated by dashed vertical lines, which are aligned to the end of the previous infection phase. The heterogeneity in infection phase duration is displayed as boxplots at the top of the graph. The expected success rate for progeny production for two virion infections (I) could not be directly measured, as virion production can only be assigned to individual cells when using very low MOI which only results in infections with one virion. Therefore, the median time to virion production in two-virion infections was based on one virion infections. The success rate of virion production was calculated assuming that each virion provided an independent chance of success. **(J)** Quantification of the time between inoculation and initiation of replication (top) or between initiation of replication and nuclear export of vRNPs (bottom) in cells infected with patient-derived IAV (left) or PR8 (right). **(K)** Success rates of vRNP replication (in infected cells) and vRNP nuclear export (in cells with replicating virus) in cells inoculated with patient-derived IAV. **(C, F, G, K)** Dots represent average values from independent experiments. Error bars represent SDs. See also Videos S4 and S5.

VIRIM-neg provides the unique opportunity to group infected cells based on the number of infecting virions, allowing us to quantify the impact of virion number per infection on the success-rate of each step in the infection cycle. This analysis revealed that multi-virion infections have higher success-rates than one-virion infections for every infection phase that was assessed (Figure 3F and 3G), with replication initiation being especially sensitive to virion number, suggesting that infections by multiple virions may enhance replication of each other (Figure 3G (top)). When summing up the success rates for all individual steps, we calculated an overall success-rate of one-virion infections of just 4% (Figure 3H), which corresponds well to the comparison in virus titer between VIRIM-neg and TCID_50_ measurements (Figure 3A). The overall success-rate of two-virion infections was increased by ∼5 fold (Figure 3I), highlighting the large impact of the number of virions per infected cell on infection success, and underscoring the importance of assessing the MOI for each infected cell when examining infection outcome. Furthermore, a major advantage of VIRIM-neg over existing live-cell techniques is that it can be applied to unmodified viruses. We leveraged this unique advantage to also map the infection cycle of patient-derived virions, which revealed a similar heterogeneity in infection cycle pathways, with distinct success-rates for individual steps in the infection cycle (Figure 3J and 3K).

### Heterogenous viral gene expression signatures underlie diverse infection pathways

The fact that the success rate of each step in viral infection is improved by co-infection of multiple virions, suggests that abortive infections are caused, at least in part, by virion-intrinsic rather than cell-intrinsic heterogeneity. Virion-intrinsic heterogeneity can include heterogeneity in vRNP content, or variable activity in transcription or replication. While vRNP packaging and replication are directly observable using VIRIM-neg, vRNP transcription is not. Therefore, to investigate whether transcriptional heterogeneity impacts infection progression, we combined VIRIM-neg and Nb^HA^ imaging with multiplexed smFISH to quantify the levels of mRNA species from all eight IAV segments, as well as three vRNA species (PB2, PB1, NP) in individual cells (Figure 4A and S6, Video S6). Multiplexed smFISH revealed remarkable differences in viral gene expression among infected cells, with anywhere between 0 and 8 different viral genome segments being transcribed, and with expression levels differing by several orders of magnitude (Figure 4B and 4C). Unsupervised clustering on the gene expression data yielded 10 distinct clusters of cells (Figure 4C, 4D, and S7A). Cluster 1 corresponds to cells lacking detectable expression of any viral genome segment, which included both uninfected cells and cells infected with IAV that did not produce any viral mRNAs (Figure 4C and S7B). Clusters 2-8 correspond to cells in which low viral mRNA levels (<100 mRNAs) were detected, with expression from only one or a few genome segments (Figure 4C), demonstrating that many infected cells transcribe a random, incomplete set of viral genome segments. The final two clusters, 9 and 10, consisted of cells with high levels of viral mRNA (Figure S7C). Cluster 9 cells have high levels of most but not all types of viral mRNAs, while most cluster 10 cells highly transcribe all eight genome segments (Figure 4C). From this analysis we conclude that in the large majority of infected cells most genome segments are transcriptionally silent.

**Figure 4.**
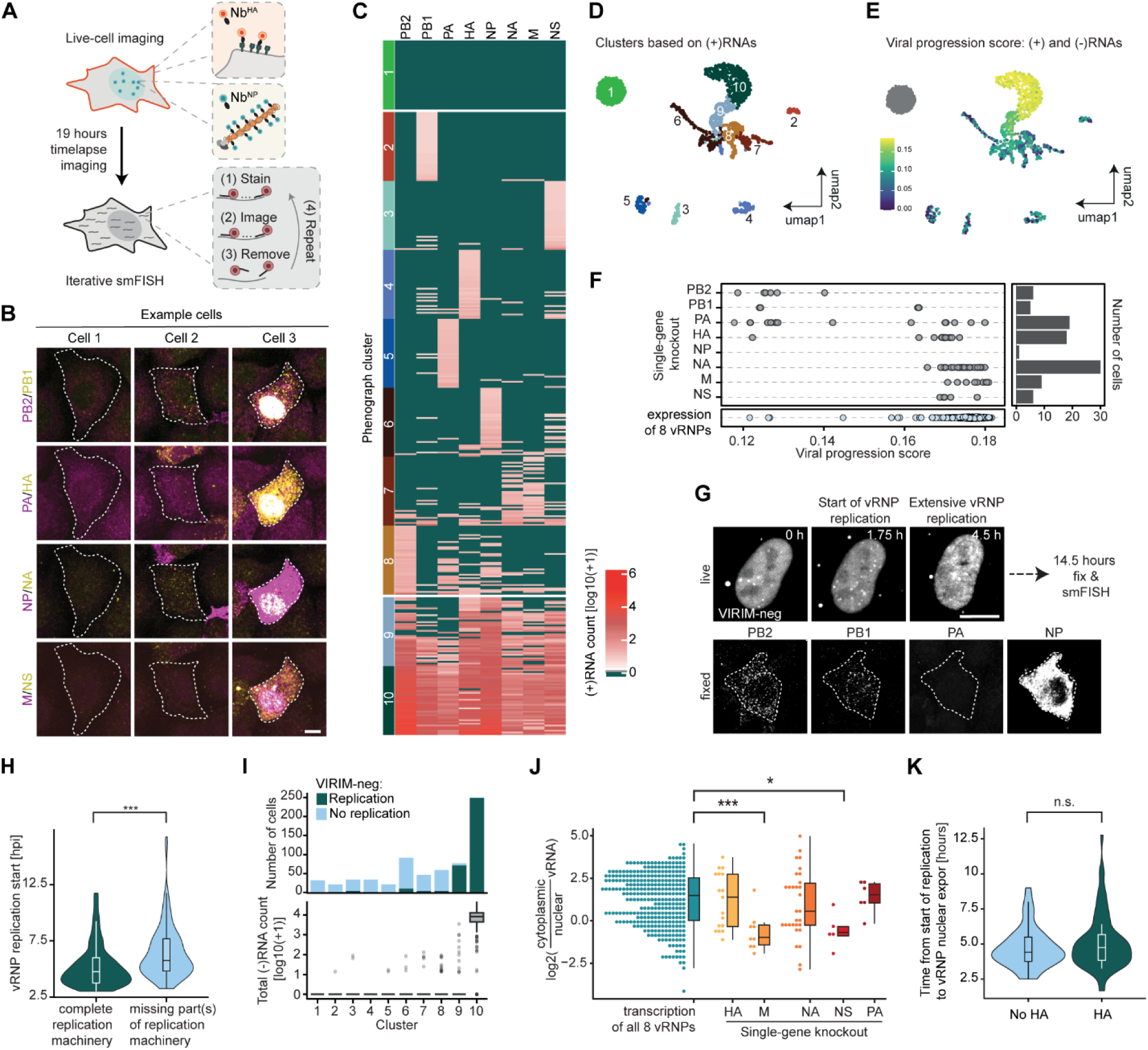
Gene expression signatures underlying heterogeneous infection progression. **(A)** Schematic of the experimental procedure of experiments combining VIRIM-neg and Nb^HA^ imaging with fixed-cell multiplexed smFISH. **(B)** Example images of multiplexed smFISH for three cells with varying levels of different viral mRNAs. Scale bar, 10 µm. **(C)** Heatmap of mRNA levels for all eight viral genome segments. Each horizontal line represents one cell. Cells are grouped based on phenograph clusters (See D and Figure S7A to S7C). 40 randomly selected cells are shown for each cluster. **(D)** UMAP projection computed on mRNA levels for all eight viral genome segments, colored according to phenograph cluster. Cluster numbers also used in (C) and (I) are shown. **(E)** The viral progression score was calculated for each cell (Figure S7D to S7F), and projected onto the UMAP representation. **(F)** Viral progression score of cells transcribing seven out of eight vRNPs, grouped based on which vRNP is transcriptionally silent. The inactive segment is indicated on the y-axis. Number of cells in each group is shown on the right. Cells for which all 8 genome segments are expressed are shown at the bottom. **(G)** Example images of live-cell VIRIM-neg (top) and smFISH of the same cell (bottom) for a cell that shows IAV replication, but lacks expression of the PA segment. Scale bars, 10 µm. **(H)** Violin plots showing the time between inoculation and first vRNP replication for cells that either express all four viral proteins needed for replication (PB1, PB2, PA and NP) or lack expression of one or more of these proteins. **(I)** Fraction of cells showing replication as assessed by VIRIM-neg (top), as well as total vRNA count (bottom) for cells in each phenograph cluster. Note that box plots are not visible for clusters 1-9 (only outliers are visible) as the majority of cells in these clusters have very low vRNA count. **(J)** Ratio between cytoplasmic and nuclear staining intensity of (-)RNAs for all single gene knockout infections. Cells were only included if they have an infection progression score that indicates that the infection is sufficiently progressed that vRNP nuclear export could occur (>0.15, see Figure S8D). **(K)** Violin plots showing the time from vRNP replication initiation to vRNP nuclear export, as determined with VIRIM-neg, for cells either expressing or lacking HA, as determined by Nb^HA^ live-cell imaging.

To understand how heterogeneous viral gene expression impacts infection progression, we computed single-cell viral progression scores, based on viral mRNA levels, as well as vRNA levels and localization, and assessed the impact of distinct viral gene expression signatures on infection progression (Figure S7D to S7G, 4E, and 4F). This approach allowed us to examine infections in which 7 of the 8 segments were transcribed, which are effectively naturally occurring ‘single-gene knockouts’, uniquely allowing assessment of the function of individual genome segments in infection progression. Studying such single-gene knockouts is difficult using genetic approaches, since most IAV genes are essential for production of virions. Analysis of single-gene knockouts revealed that a lack of expression of the segments that are not essential for replication (HA, NA, M, and NS) had only a minor effect on the viral progression score (Figure 4F). Unsurprisingly, infections lacking expression of NP, PB2, PB1, or PA typically progress less far in infection (Figure 4F). The total number of infected cells in which all segments were transcribed except either NP, PB2 or PB1 was low, likely because infections lacking one of these segments arrest during very early infection, even before the other segments are transcribed sufficiently for detection (>∼10 transcripts). Remarkably though, some PB1, PB2 and especially PA single-gene knockout cells could be identified with a relatively high viral progression score (Figure 4F). Consistently, some cells that were devoid of one of the polymerase subunit mRNAs showed unambiguous vRNP replication by VIRIM-neg (Figure 4G and 4H), indicating that (limited) genome replication can occur in the absence of expression of subunits of the replication machinery. Infected cells lacking polymerase subunit mRNAs were highly enriched in cluster 9 (Figure 4I, and S8A to S8C). These cells often showed limited levels of replication, revealing that lack of *de novo* synthesis of the viral polymerase complex results in a block of viral progression after initial replication (Figure 4I, bottom panel). These results are surprising, as previous work suggested that vRNP replication requires a second copy of the polymerase complex, likely created through *de novo* viral protein synthesis [19, 20]. Limited replication as observed by VIRIM-neg does not exclusively reflect vRNA-to-cRNA replication, since vRNA amplification was also observed in cells lacking replication subunits (Figure S8D). Possibly, the infecting virion brings along additional polymerase protein copies to drive initial rounds of replication, which may help kickstart the viral replication cycle.

We similarly assessed which viral genes play a role in vRNP nuclear export. We examined infected cells with a viral progression score of >0.15, the score range at which vRNP nuclear export typically occurs (Figure S8E). Since insufficient numbers of infected cells (<5) lacking PB2, PB1 and NP reach this stage of viral infection progression – likely because of reduced replication efficiency – these viral genes were excluded from this analysis. A lack of M or NS expression impaired vRNP export, consistent with previous reports [21-24], while a lack of HA, NA, or PA did not significantly affect vRNP export (Figure 4J). Interestingly, our finding that HA is not important for vRNP nuclear export contradicts earlier reports showing that HA buildup at the plasma membrane is needed to trigger vRNP nuclear export [25]. To confirm our findings, we also examined the timing of vRNP nuclear export in infected cells with or without HA expression, but found that vRNP nuclear export occurred at similar times (Figure 4K). To exclude that HA at cell internal membranes (which is not detected by Nb^HA^) triggered vRNP nuclear export, we examined HA mRNA expression and Nb^HA^ staining in the same cells. This analysis revealed that almost all cells lacking cell surface HA also lacked HA mRNA and thus did not contain cell internal HA protein either (Figure S8F), confirming that HA protein is not required to trigger vRNP nuclear export. In summary, these results show that the large majority of infected cells show viral gene expression defects, and that distinct gene expression signatures underly different types of infection progression failures.

### Mechanisms of gene expression heterogeneity

We next investigated the mechanisms underlying heterogeneous gene expression signatures. Absence of mRNA expression could be due to missing genome segments, which, in turn, could be caused either by failures in vRNP packaging [3, 26, 27], or, potentially, by degradation of vRNPs during infection [28]. Analysis of cells without viral replication revealed that vRNP number remained constant over time, suggesting that vRNP degradation is rare and thus likely does not contribute to the observed absence of vRNPs (Figure 5A). To assess the extent to which genome segment packaging defects could explain the observed lack of viral mRNA expression in infected cells, we quantified the frequency of missing genome segments in individual virions by smFISH. Genome segments were absent from virions at frequencies, ranging from 8%-27% (Figure 5B and 5C), consistent with previous reports [3, 26, 27]. Based on our smFISH measurements we calculated that virions contain on average ∼7 genome segments, a value very similar to the average number of vRNPs per cell as determined by VIRIM-neg (6.7 vRNPs), confirming that VIRIM-neg spot count accurately reflects genome segment number per virion. To assess whether missing segments are randomly distributed among virions, we compared the expected number of vRNPs per virion assuming that missing genome segments were randomly distributed over virions (i.e. vRNP number per virion follows a Poisson distribution) with the actual number of vRNPs per virion as determined by VIRIM-neg. We observe a broader distribution in vRNP number in the experimental data and an increased frequency of correctly packaged virions (8 vRNPs) (Figure 5D), explaining the origin of the highly diverse vRNP number per infected cell, which we found both for PR8 but also in patient-derived virions (Figure S9A). These results further suggest that vRNP packaging occurs in a cooperative manner, in line with our finding of cooperative disassembly of the vRNP bundle during vRNP entry. Cooperative packing and unpacking of vRNP bundles might be explained by vRNP-vRNP interactions within a vRNP bundle that provide increasing bundle stability with increasing numbers of vRNPs in the bundle [29, 30]. Besides packaging defects as a cause of missing genome segments, long-term time-lapse imaging of infection revealed that genome segments can also be lost during host cell division (Figure 5E). Quantification of vRNP numbers showed that both daughter cells after cell division receive approximately half of the vRNPs originally present in the mother cell (Figure 5F), indicating that vRNPs are segregated randomly during mitosis, creating two host cells with ‘viral aneuploidy’. Viral aneuploidy was especially prevalent when replication had not yet occurred at the moment of host cell division (Figure 5G). Taken together, these results indicate that missing genome segments, caused by either packaging defects or vRNP segregation during mitosis, contribute to gene expression failure.

**Fig. 5.**
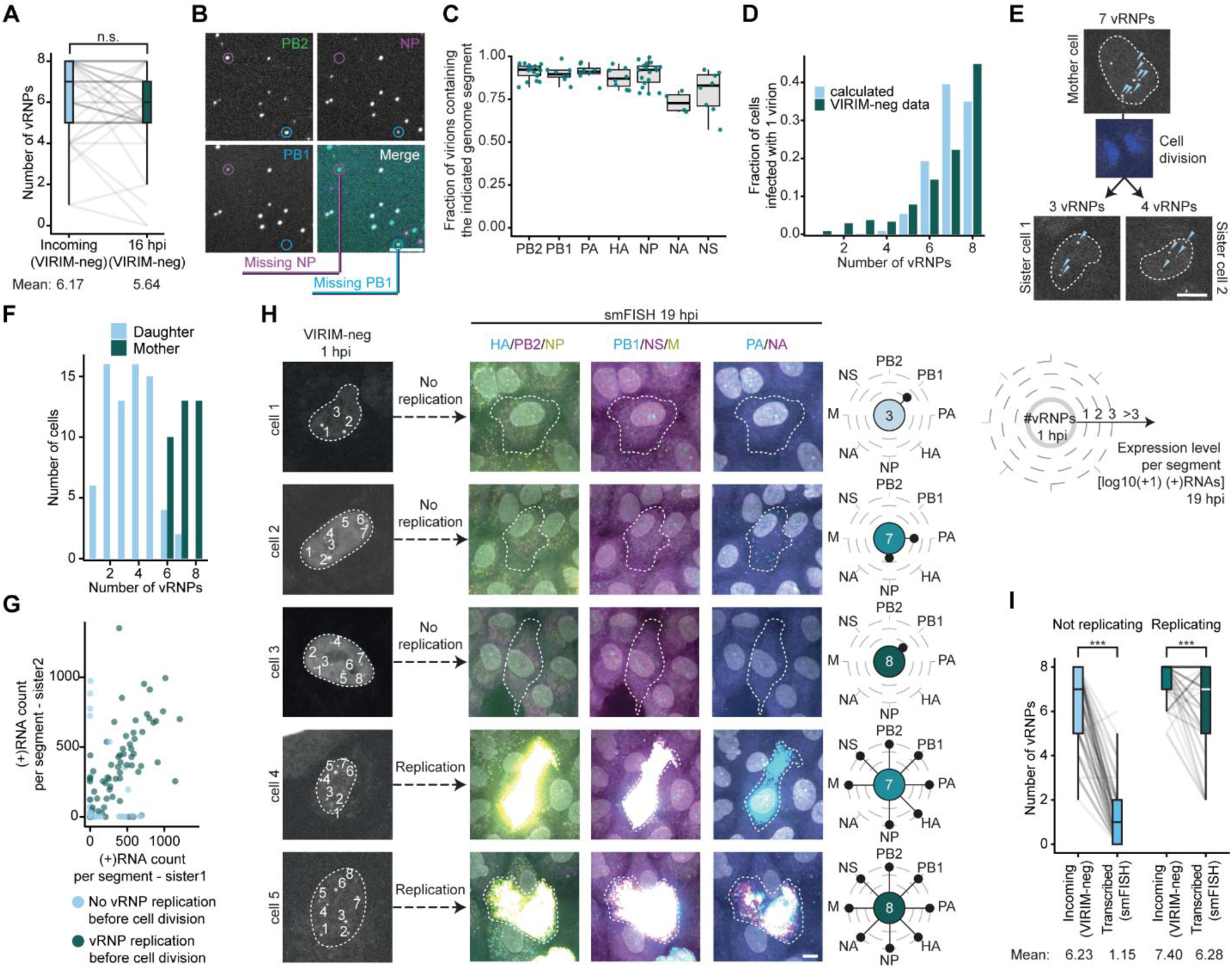
Mechanisms of viral gene expression heterogeneity. **(A)** Box plots showing the number of vRNPs per cell, as assessed by VIRIM-neg, for cells infected by a single virion in which no vRNP replication is observed. vRNP number was assessed for the same cells at both 1 and 16 hpi. The lines between box plots link individual cells in both analyses. Slight variations in spot number at early and late timepoints are likely due to technical noise in measurements. **(B-C)** IAV virions were spotted on glass and vRNAs were detected by smFISH. Representative images (B) and quantification (C) is shown. Note that genome segment M is missing from this analysis due to low signal of smFISH probes. Scale bar, 5 µm. **(D)** The number of vRNPs per virion determined experimentally by VIRIM-neg is shown (dark blue bars). Predicted distribution based on the frequency that each segment was absent from virions and assuming that missing segments are randomly distributed over virions is also shown (light blue bars). **(E)** Example images of a VIRIM-neg movie for a cell undergoing cell division. DNA is stained with Hoechst. **(F)** Quantification of the number of vRNPs per cell before and after cell division. **(G)** Comparison of mRNA expression levels between two sister cells after cell division. Each dot represents the expression levels for a single genome segment. **(H)** Example images of VIRIM-neg at 1 hpi and smFISH at 19 hpi for the same cells. Plots on the right show the number of vRNPs at 1 hpi in the center and mRNA expression levels at 19 hpi for each segment in the corresponding cell, shown as circular lollipop chart. Scale bar, 10 µm. **(I)** Box plots showing the number of vRNPs per cell, as assessed by VIRIM-neg, for cells infected by a single virion at 1 hpi (Incoming), either for cells that do or do not show replication. The number vRNPs that show mRNA expression (Transcribed) is shown for the same cells. The lines between box plots link individual cells in both analyses.

As described above, infected cells often lack genome segments, containing an average of ∼7 out of 8 genome segments. However, many infected cells express <2 genome segments, suggesting that defects other than missing genome segments contribute to the lack of viral gene expression (see Figure 4C). To uncover the origin of the discrepancy between vRNP number and number of expressed mRNAs, we compared the number of incoming vRNPs with the number of transcribed mRNAs in the same cells, for cells with or without replication. This analysis showed that many genome segments were transcriptionally silent, especially in cells without vRNP replication (Figure 5H, 5I, and S9B). Together, our results show that defective viral gene expression underlies most instances of abortive infection, and that the resulting diversity in gene expression signatures underlie heterogeneous infection cycle pathways. Furthermore, we find that defective viral gene expression is predominantly caused by transcriptional defects, and to a lesser extent due to missing vRNPs, which is caused either by packaging defects or due to unequal segregation of vRNPs during host cell division.

We finally asked whether defects in viral gene expression affect the host cell antiviral response. Typically, only a small fraction of host cells induces expression of antiviral genes (e.g. interferons) in response to IAV infection [1, 31-33]. Previous work identified the viral protein NS1 as the main antagonist of the antiviral response [1, 31-33], but how expression levels of NS1, as well as that of other viral genes, relate to antiviral response activation in infection of wildtype IAV has not been resolved. We found that IFN-λ, an antiviral gene that is upregulated in response to viral infection [34, 35], was exclusively expressed in cells with high viral mRNA expression (>100 total viral mRNAs) (Figure S10A and S10B), consistent with a recent report that viral load correlates positively with antiviral gene expression [33]. Zooming in on infected cells with at least 100 viral mRNAs, we found that only 7% of these infected cells had activated IFN-λ transcription (Figure S10B), demonstrating highly efficient suppression of antiviral gene expression by wildtype IAV. Interestingly, we found that almost all IFN-λ expressing host cells showed low or undetectable NS segment expression (Figure S10C), demonstrating that NS expression failure was a pre-requisite for host innate immune activation during a natural infection. A similar effect was not apparent for PB2 mRNA expression, despite the fact that PB2 has also been implicated as potential immune antagonists (Figure S10C) [36-38]. Lack of NS expression alone was, however, insufficient to induce IFN-λ expression, even in cells with >100 mRNAs expressed (Figure S10C, lower left quadrant). VIRIM-neg imaging showed that a lack of IFN-λ expression in NS-negative cells typically coincided with lower rates of viral genome replication (Figure S10D). These data suggest that extensive replication is another prerequisite to activate antiviral responses. Finally, we found in our live-cell imaging data that cells undergoing mitosis within several hours of infection, often showed one daughter cell with high IFN-λ expression (Figure S10E and S10F), which coincided with an absence of NS expression, indicating that the NS segment was lost in that daughter cell due to viral aneuploidy induced by host cell division. Thus, viral aneuploidy caused by mitosis could potentially affect the spread of IAV infections by inducing tissue-wide antiviral responses. Together, these data reveal how replication dynamics and viral gene expression defects together shape antiviral response activation, and identify viral aneuploidy induced by host cell division as an additional trigger for antiviral response activation.

In this study, we generate the first comprehensive single-cell kinetic map of a viral infection cycle through single vRNP and virion imaging, combined with viral transcriptomics in the same cells. We find that the vast majority of infections (∼96%) do not follow the canonical infection pathway, but undergo alternative, abortive infection pathways, and show that diverse viral gene expression signatures underlying alternative infection pathways. We identify three major pathways that cause heterogeneous viral gene expression; vRNP packaging defects, defective transcriptional activation of vRNPs and viral aneuploidy caused by host cell division. Furthermore, loss of expression of individual viral genes provided the unique opportunity to study viral gene function, including essential viral genes that cannot be knocked out genetically. This comprehensive infection map will provide a benchmark for studies detailing other IAV variants with varying pathogenicity and host range, as well as viral mutants, and clinical isolates. As such, our data will help to understand how viral evolution affects infection phenotypes and how host range and cell tropism are affected by viral genotypes, critical parameters to understand viral evolution, spread and pathogenicity.

## Acknowledgments

J.S., H.H.R, M.J.D.B., and M.E.T. are supported by a grant from the European Union (ERC, VirIm, 101044794). M.M., and M.E.T. were financially supported by the Oncode Institute, which is funded in part by the Dutch Cancer Society (KWF). T.B. and R.F. are supported by the NIAID/NIH contract HHSN272201400008C and EU4Health grant DURABLE (no. 101102733).

## Author contributions

Conceptualization, H.H.R., J.S., M.M., and M.E.T.; Data curation, H.H.R., J.S., J.P., M.M., and M.J.D.B.; Formal analysis, H.H.R., J.S., M.M., M.J.D.B., J.P., and M.E.T.; Funding acquisition, H.C., R.A.M.F., and M.E.T.; Investigation, H.H.R., J.S., M.M., J.P. and A.F.M.D.; Methodology, H.H.R., J.S., M.M., M.J.D.B., A.F.M.D., T.M.B., H.C., R.A.M.F., and M.E.T.; Project administration, H.H.R., J.S., M.M., M.J.D.B., H.C., R.A.M.F., and M.E.T.; Resources, M.M., M.J.D.B., A.F.M.D., T.M.B., H.C., R.A.M.F., and M.E.T.; Software, M.M., and M.J.D.B.; Supervision, H.H.R., H.C., R.A.M.F., and M.E.T.; Validation, H.H.R., J.S., and M.M.; Visualization, H.H.R., J.S., M.M., M.J.D.B., and M.E.T.; Writing – original draft, H.H.R., J.S., M.M., and M.E.T.; Writing – review & editing, H.H.R,. J.S., M.M., A.F.M.D., T.M.B., H.C., R.A.M.F., and M.E.T.m

## Materials and Methods

### Cell lines

A549 (ATCC, Cat# CCL-185), MDCKII and HEK293T cell lines were grown in DMEM (4.5 g/L glucose, GIBCO, Cat# 31966021) supplemented with 10% fetal bovine serum (FBS, Sigma-Aldrich, Cat# F7524) and 1% penicillin/streptomycin (GIBCO, Cat# 15140122) and regularly passaged when reaching 80-90% confluency using TryplE (GIBCO, Cat# 12605010). All cells were cultured with 5% CO_2_ at 37 °C. Cell lines were confirmed to be mycoplasma negative.

### IAV production

Generation of A/Puerto Rico/8/1934 (PR8) by a reverse genetic system was conducted as reported previously [39]. Briefly, eight plasmids encoding the individual genome segments under control of bidirectional promoters were transiently transfected into HEK293T cells using the calcium phosphate transfection method. Medium was replaced after 16 h with fresh DMEM supplemented with 2% FBS and TPCK-treated trypsin (1 µg/mL, Merck, Cat# 20233). 48 h after medium replacement, HEK293T supernatant containing virus was harvested and added to MDCKII cells for 2 h. Cells were then washed and kept in OptiMEM (Thermo Fisher Scientific, Cat# 11058021) supplemented with trypsin. MDCKII supernatant was collected after 2 to 3 days, when cytopathic effects (CPE) were observed, and stored at -80 °C. Virus stocks were used for passaging on MDCKII (unless indicated otherwise) cells at low multiplicity of infection for a maximum of two passages. To produce IAV stocks in chicken eggs, eleven-day old embryonated chicken eggs were inoculated with PR8 influenza virus at low MOI. Allantoic fluid was harvested two days post-inoculation and centrifuged for ten minutes at 2,100 *g* to remove cellular debris.

### Clinical IAV samples

Clinical IAV samples were derived from hospitalized patients that were admitted with respiratory complaints and that tested positive for IAV. Throat or nasal swaps were collected in Universal Transport Medium (Copan) according to the according to manufacturer’s protocol and stored at -80 °C until further usage.

### Design of Nb expression constructs

The VIRIM-neg imaging technique makes use of a stably expressed NP-specific nanobody (Nb^NP^) that was generated previously [8]. For visualization, Nb^NP^ was fused at its C-terminus to the bright green fluorescent protein AausFP1 [40], and a nuclear localization sequence (NLS) was added to the C-terminus of the Nb^NP^-GFP fusion protein to ensure nuclear localization. The A549 cell line expressing Nb^NP^ (A549-Nb^NP^) was used for all experiments unless stated otherwise. To quantify cytoplasmic vRNPs we generated the cell line A549-Nb^NP^cyto, in which the Nb^NP^ was fused to the green fluorophore mStaygold without NLS [41]. To label HA protein in a live cell setting, we used a previously reported nanobody that targets HA (Nb^HA^, previously described as Nb-B6) [42]. Nb^HA^ was tagged with a C-terminal superfolderGFP or mCherry and an N-terminal CD5 secretion sequence [43]. The fusion protein was expressed stably in HEK293T cells (HEK293T-Nb^HA^).

### Generation of stable cell lines

Stable cell lines were generated via lentiviral transduction as described previously [10]. In brief, the pHR lentiviral plasmid encoding the transgene of interest was co-transfected along with the packaging vectors psPax and pMD2.G using Polyethylenimine (Polysciences Inc, Cat# 23966). Transfection medium was replaced with DMEM 1 day after transfection and lentivirus collected 2 days later. Lentivirus was added to recipient A549 cells (for Nb^NP^ expression) or HEK293T cells (for Nb^HA^ expression) together with Polybrene (10 mg/mL, Santa Cruz Biotechnology Inc, Cat# TR-1003-G). Spin-infection was performed for 120 min at 2000 rpm at 25 °C. HEK293T cells expressing Nb^HA^ were selected by fluorescence-activated cell sorting (FACS) for highly expressing cells. To generate monoclonal A549-Nb^NP^ cells, single cells with low expression of Nb^NP^-GFP were sorted by FACS into 96-well plates. Low expression of Nb^NP^ is crucial to allow observation of VIRIM-neg spots over the background fluorescence of Nb^NP^-GFP. Single-cell clones were subsequently screened for low, homogenous and nuclear Nb^NP^ expression by fluorescence microscopy. An additional screening round was performed to ensure VIRIM-neg spots could be observed after infection with IAV.

To visualize IAV cytoplasmic entry (Figure 2B to 2E), a high-expressing A549-Nb^NP^ clonal cell line was selected by FACS. Due to the higher level of Nb^NP^ expression, sufficient Nb^NP^ was present in the cytoplasm to allow vRNP visualization in the cytoplasm.

### Human airway epithelial cell culture and imaging

A human airway organoid (AO) line (9209N) from an established organoid biobank was used in this study, with ethical approval granted by the NKI Institutional Review Board (IRB, M18ORG/CFMPB582) [44]. AO maintenance protocols have been described previously [18].

To generate AOs stably expressing Nb^NP^-GFP, single-cell suspensions of passage 4 organoids were infected with a lentivirus carrying the construct encoding for Nb^NP^-GFP, according to the protocol outlined previously [18]. After two months of expansion, organoids were dissociated into single cells and cells expressing low levels of GFP were isolated via fluorescence-activated cell sorting (FACS). The sorted cells were plated at high density (approximately 1300 cells/μL) in organoid culture conditions. To support cell survival, 50% conditioned media from healthy AO cultures and 5 nM heregulin beta-1 (PeproTech, Cat# 100-03) were added during the first week post-sorting [45]. The organoids were subsequently expanded using standard AO media.

For influenza A virus (IAV) infection studies, organoids were plated onto glass-bottom imaging plates. To promote 2D cell attachment, the plates were pre-coated either with 5% Cultrex Basement Membrane Extract, PathClear (R&D systems, Cat# 3432-010-01) in AdDMEM/F12 (incubated at 37°C for 30 minutes) or with 5 μg/mL invasin in PBS (incubated overnight at 4°C) [46]. Following coating, AO cells were seeded into 96-well plates (1 million cells/well) or 384-well plates (200,000 cells/well) in AO media. Media was refreshed every 3-4 days, and cells were expanded in 2D cultures for approximately two weeks prior to IAV infection experiments.

### Nb^HA^ harvesting

HEK293T-Nb^HA^ cells were grown to 95% confluency. During cell growth, Nb^HA^ accumulated in the supernatant, which was harvested at every passage (twice a week). Supernatant was purified with a 0.22 µm syringe filter to remove cell debris and stored at 4 °C until further use.

### IAV growth curve analyses

To assess the impact of Nb^NP^ expression on virus fitness, IAV growth was compared between A549 WT cells and A549-Nb^NP^ cells. Cells were grown to 70-90% confluency in 24-well plates. Cells were inoculated for 2 h with IAV (MOI=0.2), washed once with PBS and further incubated in OptiMEM supplemented with trypsin (1 µg/mL). At the indicated time points, both cell supernatant and cells were harvested and frozen at -20 °C. The concentration of infectious virions in the supernatant was quantified by a TCID_50_ assay. Intracellular levels of viral RNA were quantified by qPCR.

### TCID_50_ assay

MDCKII cells were seeded in 96-wells plates. The next day, cells were washed once in PBS, after which OptiMEM supplemented with trypsin (1 µg/mL) was added. The virus stock solutions were diluted in a 10-fold serial dilution and added to the cells. After incubation for 48 h at 37 °C, CPE was read out and virus titers were calculated according to the Spearman Kärber method [47].

### qPCR analysis of intracellular viral RNA

RNA was extracted from cells using the NucleoSpin RNA isolation kit (Machery-Nagel, Cat# 740955.50), according to manufacturer’s protocol. cDNA was synthesized with the Tetro Reverse transcriptase (bioline, Cat# BIO-65050), according to manufacturer’s protocol, using random hexamers (Thermo Fisher, Cat# SO142). qPCR was performed using IQ supermix (biorad, Cat# 1708885), with primers targeting the IAV M segment (Fw: CTTCTAACCGAGGTCGAAACG, Rv: AGGGCATTTTGGACAAAKCGTCTA) [48]. As an internal control, primers targeting the host cellular GAPDH gene were used (Fw: CACCGTCAAGGCTGAGAACGGG, Rv: GGTGAAGACGCCAGTGGACTCC). ΔΔCt values were used to calculate the fold increase of viral RNA relative to the first time point (2 h).

### Drug treatments of cells

For visualization of the nucleus during live-cell imaging experiments, DNA was stained with Hoechst 33342 (Thermo Fisher Scientific, Cat# 62249) at a final concentration of 1 µg/mL. The vacuolar-type H^+^-ATPases inhibitor bafilomycin A1 (BafA1, Merck, Cat# SML1661-.1ML) was used to block IAV cell entry. BafA1 was added at 50 nM final concentration after 1 h of inoculation, unless indicated otherwise. For experiments that include Nb^HA^ imaging, Leibovitz’s L15 medium (GIBCO, Cat# 21083027), supplemented with 10% FBS and 1% penicillin/streptomycin, was used in a 2:1 ratio with Nb^HA^-containing supernatant (see the ‘*Nb^HA^ harvesting*’ section) and added to the cells. To synchronize cells in the cell cycle, cells were arrested in G2 using the CDK1 inhibitor RO3306 (Merck, Cat# 217699-5MG) at a final concentration of 5 µM for 16 h (Figure 5). To release cells from the cell cycle block, the inhibitor was removed by washing cells 3 times with L15 medium for 5 min at 37 °C. Cells were directly used for imaging and only cells which underwent mitosis were included in the analysis.

### Cell culture for imaging

Cells were grown in a 96-well or 384-well glass-bottom plate (µ-Plate, Ibidi, Cat# 89627, or Matriplates, Brooks, Cat# MGB096-1-2-LG-L, Cat# MGB101-1-2-LG-L) such that cells were 70-90% confluent at the start of imaging experiments. Live cells were kept in L15 medium supplemented with 10% FBS and 1% penicillin/streptomycin during imaging experiments. Before live-cell imaging was initiated, cells were inoculated with virus for 1 h, after which BafA1 was added to prevent further infections, unless stated otherwise. Live-cell imaging was performed at 37 °C at atmospheric CO_2_ levels. Fixed-cell imaging was performed at r.t..

For experiments focusing on vRNP entry dynamics (Figure 2), entry into host cells was synchronized using a pH change. Briefly, cells in DMEM were pre-incubated at atmospheric CO_2_ levels for 1 h, during which time pH levels rise to ∼9.5-10. Then, virus inoculation was performed and virions were allowed to attach to cells and undergo endocytosis for 30 min. DMEM was then exchanged with L15 medium, which has a pH value in the physiological range (pH 7.0-7.4) at atmospheric CO_2_ levels and promotes rapid endosomal release of the virus. Image acquisition was started immediately after the medium exchange.

### smFISH

smFISH experiments were conducted as reported previously [10, 49, 50]. For all target RNAs, complementary, short DNA oligonucleotides (20 nucleotides long, at least 48 probes per RNA) were designed using Stellaris probe designer (https://www.biosearchtech.com/support/tools/design-software/stellaris-probe-designer) and ordered from Integrated DNA Technologies (IDT). All smFISH probes targeting a single RNA were pooled (‘smFISH probe sets’) to a concentration of 200 µM. Probe sets were subsequently labelled using fluorescent Amino-11-ddUTP (Lumiprobe, Cat# A5040), which was conjugated to Dylight-488 (Thermo Scientific, Cat# 11849370), Atto-565 or Atto-633 NHS ester (AttoTec, Cat# 72464-1MG-F and Cat# AD 633-31). Ligation of fluorescent Amino-11-ddUTP to the probes was performed using Terminal deoxynucleotidyl Transferase (TdT) (Thermo Scientific, Cat# EP0162) according to manufacturer’s protocol. After labeling, probe sets were washed 3 times in 70% ethanol and resuspended in nuclease-free water.

For smFISH staining, cells were fixed in 4% paraformaldehyde (VWR, Cat# 43368.9M) in PBS for 15 min, and subsequently permeabilized in ethanol at 4 °C for 30 min. Then, cells were washed twice in smFISH wash buffer (10% formamide (Thermo Scientific, Cat# AM9342), 2xSSC in nuclease-free water). Probe sets were diluted to 10 nM final concentration in hybridization buffer (10% formamide, 2xSSC, 10% dextran sulphate (Merck, Cat# D8906-50G) tRNA (1 mg/mL, Merck, Cat# R1753-500UN), Ribonucleoside Vanadyl Complex (2 mM, NEB, Cat# S1402S), BSA (200 µg/mL, Thermo Scientific, Cat# 240400100) in nuclease-free water). Diluted probe sets in hybridization buffer were added to cells, and hybridization was performed overnight at 37 °C. Samples were then washed twice with smFISH wash buffer supplemented with DAPI (10 µg/mL, Merck, Cat# 62248) for 1 h at 37 °C, and stored in PBS at 4 °C until imaging. For iterative smFISH staining, smFISH probes were removed through incubation of samples with 80% formamide at r.t. for 6 h. After stripping away the smFISH probes, the staining protocol was repeated for a new set of smFISH probes. Efficiency of probe removal was confirmed by imaging.

### Virion smFISH

For experiments in which colocalization between vRNA and Nb^HA^ was assessed (Figure S1F and S1G), virus stocks were diluted in Nb^HA^-containing supernatant (see *‘Nb^HA^ harvesting’*). To attach virions to the glass for imaging, the solution containing the virus was added to 96-well glass-bottom plates and incubated for 3 h at 37 °C. Subsequently, samples were fixed and stained as described in the *‘smFISH’* section. All viral genome segments were stained with probe sets labelled with Atto-633. Single z-planes were imaged by confocal microscopy. Spots in each channel were identified, and colocalization assessed in automated fashion using the ComDet FIJI plugin (https://github.com/UU-cellbiology/ComDet).

For experiments to determine the frequency that each genome segment was missing from virions (Figure 5B and 5C), virions were diluted (1:30) in PBS and added to 96-well glass-bottom plates for 3 h at 37 °C. Virions were then fixed and stained as described in the *‘smFISH’* section. Three genome segments were stained in each sample using probe sets labeled with Dylight-488, Atto-565, and Atto-633. Single z-planes were imaged by confocal microscopy. Spots in each channel were identified, and colocalization assessed in automated fashion using the ComDet FIJI plugin. Colocalization of at least two probe sets was used to identify the location of a virion. Colocalization of the third channel (either Atto-565 or Atto-633) was quantified, and reports on the fraction of virions in which the vRNA of interest is present. To assess probe set sensitivity, samples were taken along for all probe sets in each experiments, in which the same genome segment was labeled with probes sets labeled with both Atto-568 and Atto-633 and colocalization was scored to determine probe sensitivity. Each value shown in Figure 5 is corrected for the probe set sensitivity. Packaging efficiency of seven of the eight genome segments could be assessed via this method. smFISH probes for the (-)M segment failed to give sufficient signal for accurate analysis. Each datapoint indicates a separate experiment, which includes five to ten images containing ∼100-200 virions each.

### Immunofluorescence (IF) staining

#### Primary antibody labelling

The previously published monoclonal human FI6 antibody targeting the IAV HA protein [51] was a kind gift of Xander de Haan. To fluorescently label the FI6 antibody with the Alexa Fluor 647, the antibody (2.5 mg/ml) was transferred to 0.1M sodium bicarbonate (pH 8.3) buffer. The Alexa Fluor 647 NHS dye (Thermo Scientific, Cat# A20006) was dissolved in DMSO and added to the FI6 antibody in a 3:1 molar ratio. The reaction was incubated for 4 h at r.t. using head-over-head rotation. The primary labelled antibody was purified using a 10 kDa MWCO Amicon Ultra Centrifugal Filter (Merck, Cat# UFC5010), following the manufacture’s protocol, and eluted in 100 µl PBS.

#### IF staining procedure

Cells were fixed in 4% PFA in PBS for 15 min, and permeabilized with 0.5% Triton-X100 (Merck, Cat# 9036-19-5) in PBS for 5 min. After fixation, block buffer was added (2% BSA, 50 nM ammonium chloride in PBS) for 30 min, followed by 45 min incubation with the primary antibody in block buffer. Samples were washed 5 times in block buffer, and subsequently incubated for 45 min with the secondary antibody and DAPI. After incubation with secondary antibodies, cells were washed once with block buffer and once with PBS. All steps were performed at r.t., and the samples were stored in PBS at 4 °C until imaging. IF for NP was performed using the mouse monoclonal anti-Influenza A virus NP antibody HB-65 (10 µg/mL, BioXcell, Cat# BE0159) in combination with secondary Alexa Fluor 647 donkey anti-mouse antibody (10 µg/mL, Invitrogen, Ref# A31571). For HA labelling, Alexa Fluor 647 labeled FI6 antibody was used (1:100 dilution). For general cell membrane staining, the sodium potassium ATPase recombinant rabbit monoclonal antibody (10 µg/mL, Invitrogen, Ref# MA5-32184) was used as primary antibody, followed by the secondary Alexa Fluor 488 goat anti-rabbit antibody (10 µg/mL, Invitrogen, Ref# A11034).

#### Combined smFISH and immunofluorescence staining

For experiments in which IF staining was combined with smFISH staining, the complete smFISH protocol was performed as described in the *‘smFISH’* section, followed by the complete IF protocol as described in the *‘IF staining procedure’* section. For the multiplexed smFISH experiments, IF staining was performed for the sodium potassium ATPase, a protein localized to the plasma membrane, to identify cell outlines that could be used for cell segmentation. Sodium potassium ATPase staining was performed in the GFP channel, which overlapped with the green fluorescence from the Nb^HA^. However, fluorescence of the Nb^HA^ had been chemically bleached by the iterative smFISH protocol.

### Microscopy

#### Hardware

Fluorescence microscopy of all live and fixed cell experiments was performed using a Nikon TI2 inverted microscope controlled by the NIS Elements software (Nikon), equipped with a CSU-X1 spinning disc (Yokagawa) and a Prime 95B sCMOS camera (Photometrics). For imaging, a 60x/1.40 NA or a 40x/1.30 NA oil-immersion objective was used. z-drift was corrected using the Nikon perfect focus system. The microscope stage was equipped with a temperature-controlled box to conduct live-cell experiments at 37°C and fixed-cell experiments at r.t.

#### Microscopy acquisition settings

In all experiments, a 50 ms exposure time was used for all laser lines. For long-term live-cell acquisitions, a large field of view (FOV) was created by computationally stitching single FOVs post acquisition (see ‘*Live-cell image analysis pipeline’* section, ‘*Stitching’* subsection). The generation of a large FOV allows robust tracking of cells over longer periods of time, even with substantial cell migration. For example, for live-cell experiments combined with multiplexed smFISH, 210 individual FOVs organized in a 15x14 x,y-grid were acquired and stitched post-acquisition. The time interval in long-term live-cell experiments running less than 24 h was set to 15 min, unless indicated otherwise. For each x,y-position, 3 z-slices (1 µm step size) were acquired. VIRIM-neg images were acquired in the GFP channel, Hoechst 33342 in the BFP channel and Nb^HA^ either in the GFP channel or the mCherry channel Nb^HA^-GFP was used for experiments shown in Figure 1J, 2I, 2J, 3F to 3I, 4 (progression score), S3A, S3F to S3H, S6A, S7D to S7G, and S8E, and Nb^HA^-mCherry as shown in Figure 2F, 3D, 3E, 4K, S3B to S3E, S5A, S5D, and S5E.

Short-term imaging for counting of the number of vRNP spots (Figure S1D, 3C, S8C, 5D, and S9A) or analyses of vRNP entry dynamics (Figure 2A to 2E) was conducted without delay between acquired images (using 50 ms exposure, the time interval between subsequent images averaged 140 ms) using a single z-slice. To determine vRNP spot number per cell, single x,y positions were imaged for 20 s. For virus entry experiments, single FOVs were imaged immediately after pH value adjustment for 2 to 10 min.

For experiments in which live-cell imaging was combined with fixed-cell imaging, fixation of cells was performed immediately after acquisition of the last time point of the live-cell experiment. After staining, either following the smFISH or IF protocol, samples were aligned to the same x,y positions acquired during the live-cell experiment. For each x,y-position, 5 to 7 z-slices with 1 µm step size (stained cells), or a single z-slice (stained virus particles) were imaged.

### Image and data analyses

#### Maximum intensity projection of z-slices

Before further analysis, maximum intensity projections of all z-slices were generated by either Nikon NIS-Elements AR (5.21.03), or upon data import using Numpy (1.23.5).

#### Automated processing of live-cell imaging data

All live-cell image analysis was performed using Python (3.9.7).

##### Illumination correction

Acquisition of images with most microscopes leads to images with non-homogeneous illumination, most commonly seen by signal decreasing towards the edges of the acquired FOV. Illumination correction for live-cell image datasets was performed using images obtained from a concentrated dye solution (4 μg/mL DyLight 488-NHS Ester), and dark images. 100 images at multiple x,y positions were acquired for both light and dark exposures, which were subsequently median-projected and used to calculate a correction matrix. The correction matrix was then applied to every single FOV.

##### Image stitching

Following illumination correction, the individual FOVs were stitched together into a large FOV. Stitching parameters were computed using the MIST algorithm in ImageJ on the timepoint halfway through the time-lapse and these stitching parameters were then applied to all other timepoints [52, 53]. To launch stitching in ImageJ from python, the PyImageJ package was used.

##### Segmentation

Segmentation of nuclei was performed on the Hoechst images, using CellPose (2.2.3) with the pre-trained ‘nuclei’ model and an average object diameter of 80 [54].

##### Tracking

Tracking of cells over time was performed using the generated segmentation mask objects and the simple LAP track mode of TrackMate (7.9.2), with settings: max frame gap = 5, linking max distance = 60, gap closing max distance = 100, enable track splitting = False, enable track merging = False [55].

##### Automated Nb^HA^ quantification

Measurement of object intensity was performed using regionprops_table from scikit-image (0.22.0). To quantify the intensity for Nb^HA^ signal, a custom function was added to the regionprops that computes the 1 percentile of all pixels in the nuclear area. The 1 percentile value was then background subtracted based on the mean signal of uninfected cells (based on Nb^NP^) for all timepoints. To integrate data from replicate (n=3) experiments, the baseline subtracted intensity values were scaled between the 1^st^ and 99^th^ percentile per replicate irrespective of timepoint.

##### Mitosis calling

To assess the interplay of viral infection, replication and host cell mitosis we developed an approach to detect mitotic events and link sister-cells in an automated manner. Detection of mitotic cells was achieved by:

1. Extracting morphological and DAPI intensity features for all cells
2. Annotating a subset of cells in interphase and mitosis. Cells were chosen randomly across the whole time-lapse
3. Training a random forest classifier on the annotated cells
4. Applying the model to the rest of the data

For the cells and timepoints in which a mitotic event was predicted by the model, we searched for tracks that were newly initiated within a certain window of space and time (e.g. newly initiated tracks within 20 pixels in x- and y and within three timepoints in time). For each mitotic event, the newly initiated tracks within this window were then extracted and used for sister-cell linking. The predicted sister-cell linkages that were used for data analysis were validated by checking the time-lapse movies manually.

##### Creating single-cell grids and animations

The single-object-grid tool (https://github.com/TanenbaumLab/SingleObjectGrid) was used to crop a small FOV around tracked cells in order to create stabilized timeseries with the nuclear centroid centered. The napari-animation plugin was used to generate video output from the napari viewer to create supplemental videos.

#### Live-cell imaging analyses

##### vRNP count and determination of the number of infecting virions per cell

To determine the number of vRNPs in the nucleus or cytoplasm, infected cells were imaged for a short period of time as described in the ‘*Microscopy acquisition settings*’. Fluorescent spots were counted manually. Only cells, which were located completely in the FOV were used for analysis of spot count. If VIRIM-neg levels were too high or too low to reliably determine the spot count, the cell was excluded from the analysis. Since the A549-Nb^NP^cyto cell line was polyclonal, some cells showed stationary background spots in the GFP channel, which could readily be discriminated from VIRIM-neg spots based on their mobility. Such cells were excluded from further analysis.

To determine the number of infecting virions within a single cell, the number of VIRIM-neg spots was scored as indicated above. If the number ranged between 1-8, the cells was assigned to the group of one-virion infections, between 9-16 as two-virion infections, and >16 as infections with more than two virions. For most experiments, 20 s streaming acquisition was performed after 1h of inoculation to score the number of infecting virions as described in ‘*Microscopy acquisition* settings’. For experiments in which long-term imaging was started simultaneously with inoculation (Figure S2C), the number of infecting virions was based on spot count in single z-slices.

##### Quantification of vRNP entry dynamics

For the quantification of the duration between pH adjustment and endosomal fusion, we determined the time from pH adjustment to the first observation of a (largely) immobile VIRIM-neg spot in the cytoplasm. For this analysis we included only spots that appeared after the start of imaging and that later split into smaller, mobile spots during the observation period. All other spots were excluded from the analysis, since such spots likely represent background spots. For analysis of the dynamics of endosomal release of vRNPs, only spots were included that remained in the z-plane that was imaged during the course of imaging. The first moment a spatial separation was observed between the bright immobile spot and a mobile spot was called as the moment of release of the genome segment.

##### Annotations of viral life cycle events in VIRIM-neg time-lapse imaging

All annotations of the life cycle stages imaged with VIRIM-neg were performed manually.

The moment of infection: The moment of infection was determined based on the first VIRIM-neg spots appearing in the nucleus. If several viral particles entered the nucleus at different time points, the moment of detecting the first spot was used.

The moment of vRNP replication: The moment of vRNP replication was determined based on the first increase in VIRIM-neg spot number after vRNP entry. The increase in number of vRNP spots had to be visible for at least three consecutive points in time of which the first one is defined as the start time of replication. In cells in which the virus replicated, single-molecule resolution was often lost after several rounds of replication, which was labeled as “extensive” replication (Figure 1F and S8A). Maintaining single-molecule resolution of vRNPs until the end of imaging despite vRNP replication indicates low-level vRNP replication, and was labeled as “limited replication” (Figure S8A). It is noteworthy that the same fluorescent spot can appear in multiple z-slices due to fast diffusion of vRNPs. Consequently, a single vRNP occasionally appears twice in images of MaxIP projection of z-slices. Therefore, an increase in spot number was determined by the trend of the average spot count per cell over time.

The moment of vRNP nuclear export: The start of vRNP nuclear export was determined based on the first detectable decrease in nuclear GFP intensity with a concomitant cytoplasmic GFP intensity increase.

##### Annotations of HA staining and progeny formation using Nb^HA^

For determination of the moment of HA appearance at the plasma membrane, manual annotations were performed. Cells were considered positive for HA if a plasma membrane outline became detectable in the Nb^HA^ channel. The first moment a plasma membrane outline became apparent was annotated as the moment of HA appearance.

Progeny release was analyzed in experiments with low MOI (MOI < 0.05) to ensure that HA-expressing cells were spatially distant from each other (at least separated by one FOV). The spatial distance between HA-positive cells facilitated the identification of the cell from which progeny virions originated, as progeny spread out over large areas. Progeny release was determined by an increasing number of Nb^HA^ spots that were spatially enriched in proximity to an HA-positive cell.

##### NP IF for the validation of the vRNP nuclear export phenotype

To validate that vRNP export could be called accurately from VIRIM-neg live-cell imaging data, cells were first imaged by VIRIM-neg, after which cells were fixed and NP was stained to examine nuclear-cytoplasmic translocation of NP (which occurs along with vRNA nuclear export). For cells in which vRNP replication occurred (as assessed by VIRIM-neg), vRNP nuclear export was called manually based on VIRIM-neg to identify cells with or without vRNP nuclear export. The NP immunofluorescence signal in nucleus and cytoplasm of both groups was analyzed in automated fashion. The cell outline segmentation was performed using an IF staining of the sodium potassium pump ATPase, which is localized to the plasma membrane. DAPI staining was used for nuclear segmentation. After background subtraction (using fluorescence levels in uninfected cells), for each cell, the ratio between cytoplasmic and nuclear NP intensities was determined.

##### Colocalization Analysis of VIRIM-neg and viral RNA smFISH

Cells were incubated with virus inoculum and imaged for 3 h, allowing initiation of vRNP replication in a subset of cells. Cell were subsequently fixed and smFISH staining was performed, as described under ‘*smFISH*’. (-)PB2 was stained using probes labeled with Atto-568, and (+)PB1 was stained using probes labeled with Atto-633. Images were acquired with a z-step size of 0.5 µm (see section ‘*Microscopy acquisition settings’*). To quantify colocalization of VIRIM-neg spots with (-)RNA smFISH, images were processed as follows; first, nuclei were segmented in 3D using the GFP signal from the nanobody. To enhance the performance of segmentation and remove the VIRIM-neg spots, the GFP images were first smoothed using a gaussian filter with sigma=2. The smoothed GFP images were then used for 3D nuclear segmentation with cellpose 2.0, using the parameters: *Choose_3D_mode = "3D predictions", Object_diameter = 80, Anisotropy = 3, Min_size = 20000, mask_threshold=0, model = models.Cellpose(gpu=False, model_type=’nuclei’)* [54]. As a second step, the VIRIM-neg and (-)RNA smFISH spots were detected using deepblink [56]. More specifically, the deepblink model ‘smfish2’ was applied with *probability = 0.000001*. The spot detection was performed on each z-slice separately. To ensure that the same spot is not detected multiple adjacent z-slices, a 3D refinement step was included to find the center of the spot in the z-dimension, as described previously (https://github.com/BBQuercus/deepBlink/blob/master/examples/3d_prediction.ipynb). In brief, detected spots are tracked along the z-axis and the z-slice with the highest intensity is selected as the z-coordinate of the spot. Thirdly, since fixation can affect the VIRIM-neg spots, cells with clear VIRIM-neg spots were manually selected. Finally, colocalization was assessed by quantifying whether a VIRIM-neg spot was found within a 3-pixel distance (550nm in xy, 1.5um in z) of the detected (-)RNA spot. Since colocalization can occur due to randomness rather than specific colocalization, especially if the number of spots is high or the volume in which spots are detected is small, we analyzed the amount of random colocalization to ensure that the observed values are significant. To determine the amount of random colocalization, the same segmented nuclear 3D masks as above were used and the same number of spots that was detected for each nucleus was seeded randomly within the nuclear volume. Colocalization was then calculated as described above and compared to the observed values.

##### Quantification of VIRIM-neg spot count and (-)RNA smFISH signal intensity after vRNP replication

To corroborate that VIRIM-neg spot number increase, as observed in time-lapse imaging, accurately reports on vRNP replication, cells were first imaged using VIRIM-neg to call replication, after which vRNA content was assessed by smFISH. Cells were inoculated with virus, and imaged for 4 h. Cells were then grouped based on the number of infecting virions (1, 2, >2), and based on whether vRNP replication was observed by VIRIM-neg, see sections ‘*Annotations of viral life cycle events in VIRIM-neg time-lapse imaging*’ and ‘*vRNP count and determination of the number of infecting virions per cell*’. For each of the resulting six groups of cells, the count of (-)RNA smFISH spots was determined.

##### Calculation of expected success rates in infections with more than one virion

The expected success rates for vRNP replication, vRNP nuclear export, HA synthesis and progeny release for two or three virions (Figure 3G) was calculated by assuming that each virion has an independent and equal chance to successfully accomplish the respective infection cycle step. Calculations for expected success of two virion infections were performed using the equation:

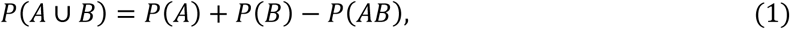

and calculations of the expected success rates of three virion infections using the equation:

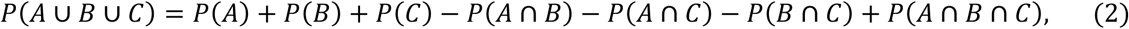

where P(*A*), P(*B*), and P(*C*) are the success rates of the individual virions for the indicated infection cycle phase, i.e. the one-virion success rates (see Figure 3F). The individual data points of the calculated success rate in Figure 3G were based on the success rate values obtained in the independent experimental repeats shown in Figure 3F.

### Analysis of experiments with multiplexed smFISH and linking to live-cell imaging (Figure 4, 5, S6, S7, S8, and S10)

#### Image analysis

##### Overview

Since live-cell data was generated in large FOVs (see ‘*Microscopy acquisition settings’*), the multiplexed smFISH readouts that were subsequently generated were also acquired in large FOVs, so that all single cells could be easily matched between live and fixed data. To achieve interpretable information from all rounds of the iterative smFISH readout (see ‘*smFISH’*) that can be matched with the live-cell imaging data, several key steps have to be performed, which are described in more detail below. All image analysis was performed using python 3.9.15. More specific packages and functions are mentioned in the separate sections below.

##### Illumination correction

To correct for non-homogeneous illumination in the multiplexed smFISH experiments, we used post-acquisition illumination correction using the BaSiC algorithm [57]. Post-acquisition illumination correction allows calculation of a correction model to correct uneven illumination, given that there are enough images to measure uneven illumination. The correction model is calculated for each channel separately, since uneven illumination can be specific for optical setups of different channels. To ensure that there are sufficient images to calculate the correction, all images that were acquired across the whole experiment (from all iterations of smFISH staining) were collected, resulting in ∼2,000 images per channel. The python implementation of the BaSiC illumination algorithm was used to compute a separate illumination correction model for each channel using the function and parameters *BaSiC (get_darkfield=True, smoothness_flatfield=1)* [57]. The models were applied to the respective channels and the corrected images were saved.

##### Image stitching

Following illumination correction, the individual FOVs were stitched together into a large FOV using the MIST algorithm in ImageJ. Stitching was performed on each staining round separately, since the stage could have shifted between iterations of smFISH. To make sure that stitching is consistent between channels of the same acquisition, the best stitching parameters were first computed on the DAPI channel (that is present in each acquisition of the iterative smFISH experiments) and these stitching parameters were then applied to all channels of the same acquisition.

##### Image registration of staining rounds

To allow for quantitative measurements of the same cells across all staining rounds, it is important that the large FOVs that were generated are aligned to each other. The image alignment, or registration, was performed on the DAPI channels of each staining round and then applied to all other channels of the same staining round. To perform registration of one image to another, a reference image has to be selected. In this case, the stitched DAPI image from a staining round half-way through all staining rounds was selected as the reference image. The reference DAPI image was then used to compute the best transformation for the stitched DAPI image of each acquisition, yielding a transformation that is specific for each staining round and is applied to the other channels of the respective staining round. Registration was performed using the python package pystackreg, a python implementation of the StackReg algorithm [58]. More specifically, the registration was performed using affine transformation of the stitched DAPI image (Figure S6A).

Since only regions which were covered by all staining iterations are of interest, all the stitched and registered images were cropped to the region which was overlapping between all iterations.

##### Nuclear and cell segmentation

After illumination correction, stitching and registration, segmentation was performed to extract quantitative measurements for all cells and nuclei across all the stainings. Segmentation of nuclei and cells was performed using cellpose 2.0 [59]. To segment nuclei, DAPI staining was used. Segmentation of nuclei was performed using the model_type=’nuclei’ with an average object diameter of 100. For cell segmentation, the membrane staining was combined with the DAPI staining to create a two-color image and cell segmentation was performed using the model_type=’cyto2’ with an average object diameter of 240.

The resulting segmentation masks were then visualized together with the images using the napari viewer and segmentation errors were manually corrected. Corrected segmentation masks were saved for further processing [60].

##### smFISH spot detection

To quantify the number of viral and host RNAs per cell, single smFISH spots have to be detected and counted. However, expression levels of viral genes in particular range over multiple orders of magnitude, yielding cells that are too full of RNA to distinguish single smFISH spots. To address this problem, dense regions – regions in which individual smFISH spots cannot be distinguished - were decomposed into single spots, as described previously [61]. Importantly, we are likely underestimating the real number of RNAs in cells with highly progressed infections because of the maximum intensity projection that is performed before starting with image analysis.

Spot detection and decomposition of dense regions yields a table with x,y-coordinates for each spot that was detected, which are used for counting spots per cell, as described in *Single-cell feature extraction*.

Since the multiplexed smFISH is generated by sequential rounds of staining, imaging and destaining, it is important that the spot counts are not affected by signal carried over from a previous round of staining into the new round. To validate this the cells were imaged after each round of destaining and the destaining rounds were processed as described above. For each destaining, the spot detection parameters of the following staining round were applied, to make sure that the same criteria for spot detection are applied as in the staining to which its signal could be carried over. We found that signal was removed very efficiently in destaining rounds (Figure S6B).

##### Single-cell feature extraction

To count the number of RNA molecules in nuclei and cells, the computed segmentation masks (as described in ‘*Nuclear and cell segmentation’*) and spot coordinates (as described in ‘*smFISH spot detection’*) are combined. The spots are counted per staining and segmentation mask, yielding a table that contains the number of detected spots per mask and staining. This procedure is performed for both nuclear and cell segmentation. To extract cytoplasmic RNA counts, the nuclear counts were subtracted from the whole-cell counts. To quantify intensity of HA^Nb^ for single cells, the regionprops function from scikit-image was used [62].

To assess whether our experimental and analysis workflow yields consistent results across staining rounds, we included one staining (PB2 negative sense staining) in round two and five and analyzed both staining rounds as described above. When comparing single-cell spot counts between the two rounds, spot counts were very consistent, with a pearson correlation of 0.95 between the two rounds (Figure S6C).

##### Spillover correction

Due to the high dynamic range of viral gene expression – ranging from a few molecules to hundreds of thousands – it is crucial that spot count quantifications for cells in which none or only a few RNAs are present, are not affected by highly positive neighboring cells. Spillover from highly positive cells can occur because of issues in stitching, registration or cell segmentation, or a combination thereof. To address this challenge, we developed a new approach to correct for this type of spillover using the following steps for every single cell and for every staining:

1. The cell segmentation is iteratively eroded by several pixels, therefore creating new segmentation in which the outermost pixels are removed, making the cell mask smaller.
2. For each of the (eroded) segmentations, the spot count and the area of the segmentation is extracted.
3. For the next steps, only segmentation masks with a defined minimal area are used, to make sure that small cells are not shrunk too much.

a. If the smallest three erosions have a spot count of zero, then set spot count to zero
b. Otherwise, calculate the corrected number of spots by performing robust linear regression on the extracted spot counts as a function of the cell area.

If a cell has a lot of spots in the outer layers of its segmentation, the first counts are high but then values drop off abruptly of the mask is eroded. The robust linear regression disregards these values because they are considered as outliers, therefore correcting the value to zero or lowly positive, depending on whether the counts are zero after the drop off – indicative of a negative cell - or if a linear scaling is observed – indicative of a lowly positive cell that was affected by spillover (Figure S6D). For cells that are not affected by spillover, the spot count should scale linearly with cell size and the count is therefore largely unaffected by the correction (Figure S6D and S6E).

##### Linking fixed cells to live cell data

To link cells between the live-cell imaging and the multiplexed smFISH experiments, nuclear segmentation masks of both the final timepoint of live-cell imaging and the multiplexed smFISH were used. Both segmentation masks were binarized. The binarized nuclear segmentation from the fixed data was then registered to the binarized nuclear segmentation of the live-cell data using registration as described in ‘*Image registration of staining rounds’*. The computed transformation was then applied to the non-binarized nuclear segmentations from the fixed data. Finally, for each nucleus in the live-cell data the most abundant pixel value (mode) of the transformed fixed nuclear segmentations was extracted, yielding a table with matched live-cell nuclear segmentation labels and fixed-cell nuclear segmentation labels which was saved for further processing.

##### Calculating RNA count cut-off values for uninfected cells

Due to noise arising from the smFISH protocol or the subsequent analysis thereof, it is important that in particular for viral RNAs no false positive cells are introduced, which would distort the analysis of infected cells. To ensure that technical noise is not affecting our smFISH readouts, we therefore took advantage of the matched VIRIM-neg information to annotate around two hundred uninfected cells based on the live-cell imaging. The uninfected cells were used to calculate the 95^th^-percentiles for spot counts in each viral smFISH staining, to make sure that we have a very low false discovery rate of viral RNAs. The 95^th^- percentiles were then used to define a lower threshold for each viral staining, below which all values were set to zero (Figure S6D).

#### Data analysis

All data analysis was performed using R version 4.2.2 and Rstudio version 2022.12.0+353 [63]. All plotting was performed using ggplot2 version 3.4.4 [64]. General data wrangling was performed using dplyr version 1.1.4 [65].

##### Phenograph clustering

Phenograph clustering was performed on the eight measured viral (+)mRNAs [66]. To determine the best k-parameter for phenograph clustering, a range of k-parameters was selected and for each, phenograph clustering was performed on the dataset. Silhouette score was then used as a measure of clustering quality, as defined by the similarity within vs between clusters. In addition to silhouette score, the resulting cluster size was taken into account and k=40 was determined as the best parameter (Figure S7A).

##### UMAP

UMAP projections were computed using the umap function from the umap package version 0.2.10.0 [67]. The umap function was run on the eight measured viral (+)mRNAs, with default umap parameters, except min_dist=0.9.

##### Viral progression score (Figure S7E)

To compute the viral progression score the following features were used:

- total viral (+)RNA in the nucleus and cytoplasm, calculated by summing up the eight viral (+)RNAs that were measured
- total viral (-)RNA in the nucleus and cytoplasm, calculated by summing up the three viral (-)RNAs that were measured: PB2, PB1 and NP
- mean HA protein intensity of the eroded cell segmentation. Cell segmentations were eroded by 30 pixels to reduce spillover from highly HA positive cells into neighboring cells.

All features were first log-transformed using y = log10(x+1) and the log-transformed features were scaled using the scale function of the base R package.

To compute the progression of infection score, cells were first reduced into 2D space using the DiffusionMap function of the destiny package with default settings except k=800 (Figure S7D) [68]. Analysis of the diffusion maps showed a group of cells that was distinct from the rest of cells. Upon closer inspection these cells turned out to be only (-)RNA positive because they were neighboring highly infected cells that were likely budding new virus that attached to neighboring cells. Since these cells are not reflecting an ongoing infection but rather proximity to a highly infected cells, we removed them from downstream analysis. Using k-means with default parameters except centers=4 and n_start=50 allowed for easy detection and removal. Upon removal of the false positive (-)RNA cells, the diffusion map was computed again on the remaining cells using the same parameters as mentioned above. To align cells along a continuous trajectory through the diffusion map, the R package of the slingshot algorithm was used (Figure S7D) [69]. The positions for single cells along the trajectory through the diffusion map were then extracted using the getCurves and slingPseudotime functions. For all subsequent analyses, the viral progression score refers to this computed trajectory, which is also shown in Figure S7D (blue arrow).

###### Statistical testing

Unless indicated otherwise, for statistical testing a two-sided t-test was performed. For paired data (Figure 5A and 5I), a paired, two-sided t-test was performed. P-values are indicated as n.s., *, **, *** for non-significant, p-value < 0.05, p-value < 0.01 and p-value < 0.001, respectively.

---

**Figure S1.**
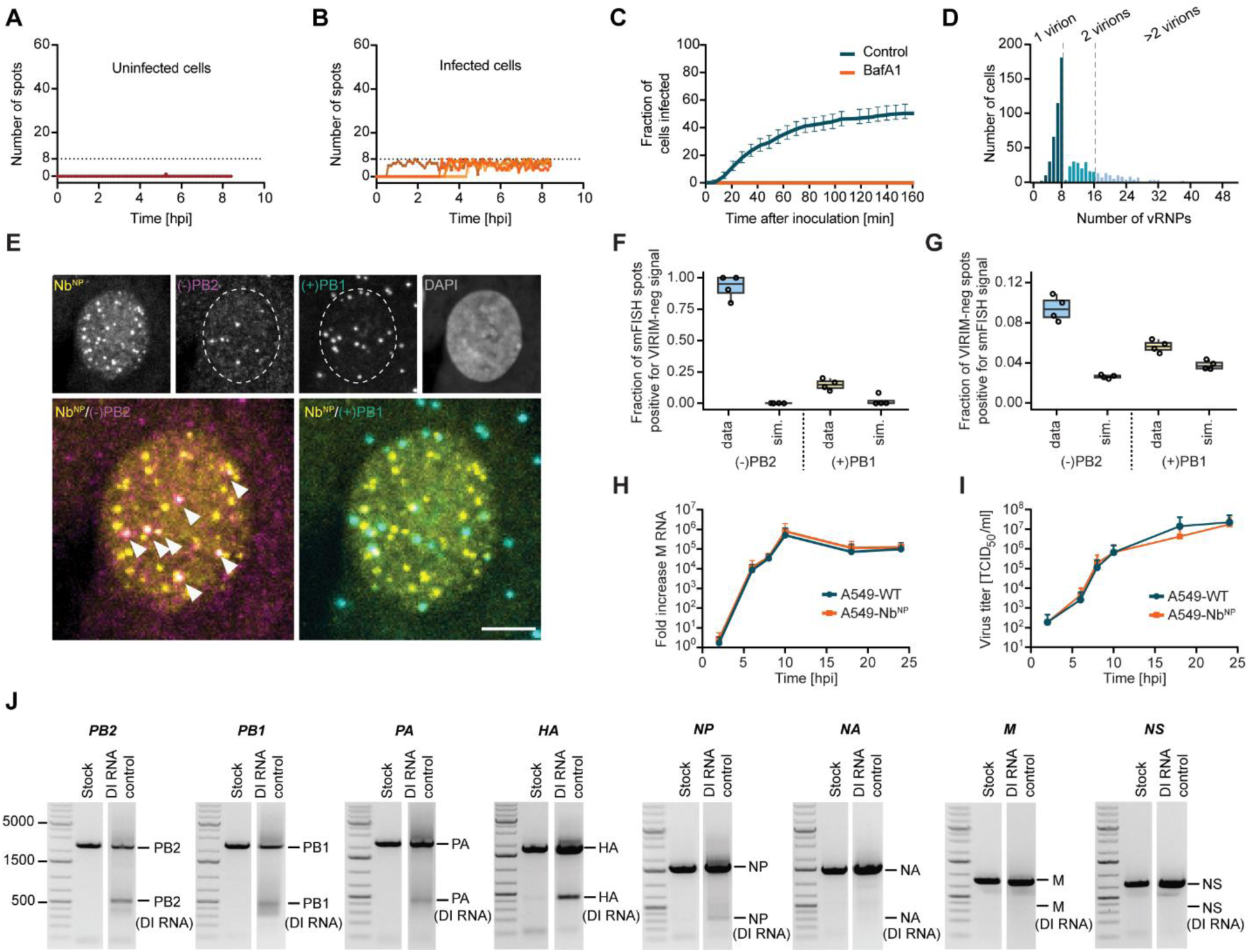
Visualization of single IAV vRNPs in live cells by VIRIM-neg. **(A-B)** Number of VIRIM-neg spots per cell over time for non-infected **(A)** or infected **(B)** cells within a population of A549-Nb^NP^ cells inoculated with IAV. Every line represents a single cell, data for four cells is shown. All traces are aligned to the start of image acquisition, which was initiated immediately following virus addition. Dashed line indicates 8 VIRIM-neg spots. **(C)** Quantification of the fraction of cells that was infected over time after inoculation with IAV, either in the presence or absence of BafA1, an inhibitor of IAV entry. Image acquisition was initiated immediately following virus addition. Error bars represent the 95% confidence interval. **(D)** Quantification of the number of vRNPs per infected cell 1 hpi. Maximal vRNP count expected for infections by a single virion (8) or by two virions (16) is indicated by dashed lines. **(E)** Example images of smFISH staining and VIRIM-neg (Nb^NP^) in A549-Nb^NP^ cells 3 hpi. Cells were stained by smFISH for (-)PB2 vRNAs (magenta) and (+)PB1 RNAs (cyan). White arrowheads indicate (-)PB2 vRNAs colocalizing with Nb^NP^ signal (yellow). Note that (+)PB1 RNAs (cyan) show little colocalization with VIRIM-neg spots, indicating that most (+)PB1 RNAs represent mRNAs rather than cRNAs. Scale bars, 5 µm. **(F)** Quantification of the colocalization between either (-)PB2, or (+)PB1, and VIRIM-neg spots of images as shown in **E**. Simulations of the expected level of random colocalization are based on the number of VIRIM-neg and smFISH spots in each nucleus (simulation (sim.)). Dots represent averages of independent experiments. **(G)** Quantification of the colocalization between VIRIM-neg spots and either (-)PB2 or (+)PB1 (exp) spots of images as shown in **E**. Simulations of the expected level of random colocalization are based on the number of VIRIM-neg and smFISH spots in each nucleus (simulation (sim.)). Dots represent averages of independent experiments. **(H,I)** Virus replication curves for the indicated cell lines as assessed by intracellular viral RNA quantification by qPCR **(H)** or infectious virus particle production as assessed by TCID_50_ assay **(I)**. Error bars indicate SD. **(J)** Agarose gel analysis of defective interfering RNAs (DI RNA). Shown are an IAV stock used for VIRIM-neg experiments (left lanes) and a positive control IAV stock created by high-MOI passaging (MOI=25) to induce formation DI RNAs (right lanes). Each of the eight genome segments were PCR amplified, and full length and smaller fragments (reflecting DI RNAs) were visualized on an agarose gel. For each genome segment, the expected full-length and DIP fragment sizes are indicated.

**Figure S2.**
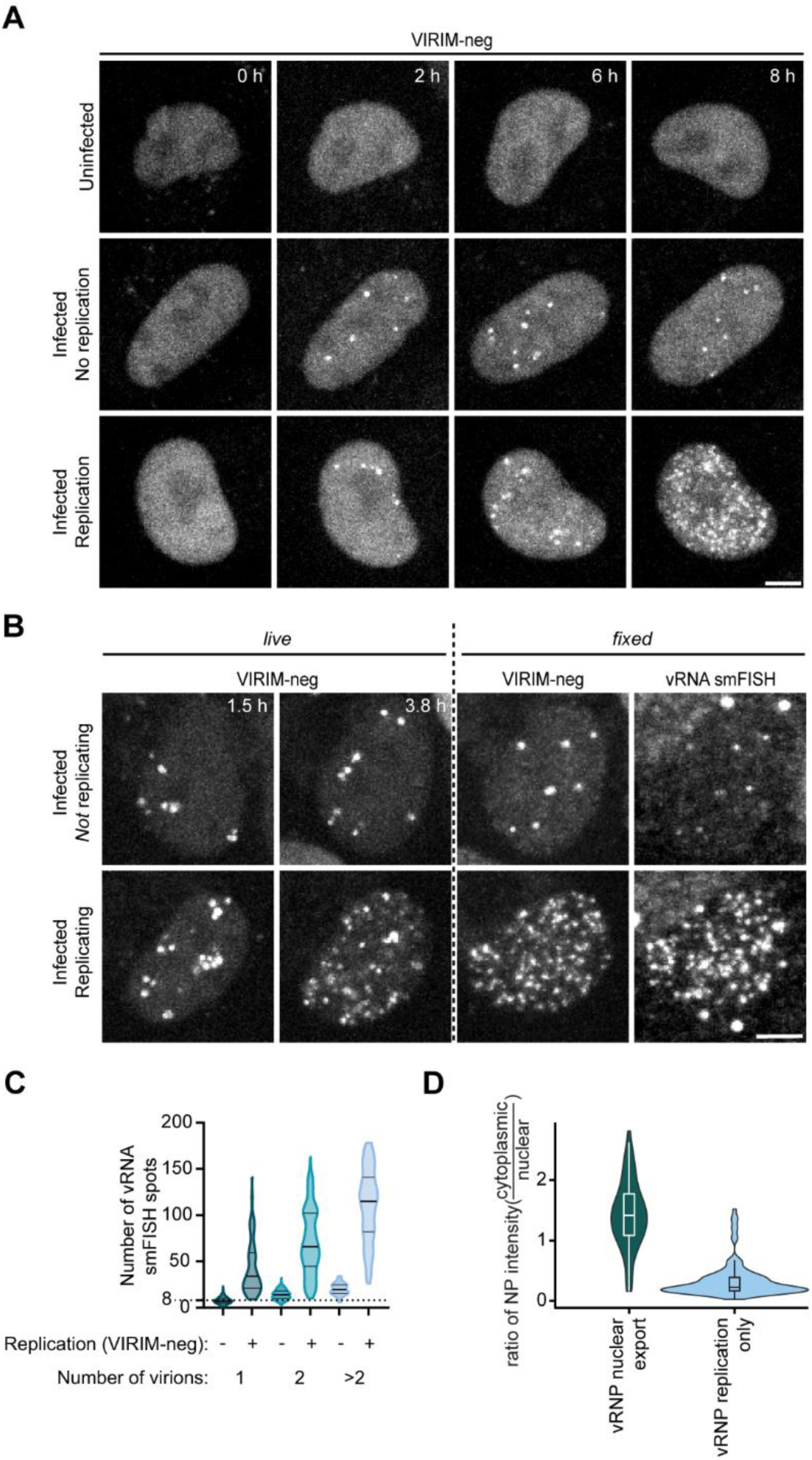
Visualization of IAV vRNP replication and nuclear export. **(A)** A549-Nb^NP^ cells were inoculated with IAV. Representative example images of time-lapse movies of an uninfected cell (top), an infected cell without vRNP replication (middle) and an infected cell with vRNP replication (bottom). **(B)** Comparison of the number of VIRIM-neg and all vRNA smFISH spots in nuclei of IAV inoculated cells, either with (bottom) or without (top) vRNP replication. 4 hpi cells were fixed and all vRNAs were stained by smFISH staining. Replication status and virion count were determined by VIRIM-neg. **(C)** Quantification of the number of VIRIM-neg and all vRNA smFISH spots in cells 4 hpi as shown in (F). vRNP replication status and virion count per cell were assessed by VIRIM-neg. **(D)** Quantification of the NP IF signal in the nucleus and cytoplasm of cells for which vRNP nuclear export either was or was not observed using VIRIM-neg. Cells were fixed 17 h post inoculation. Scale bars, 5 µm.

**Figure S3.**
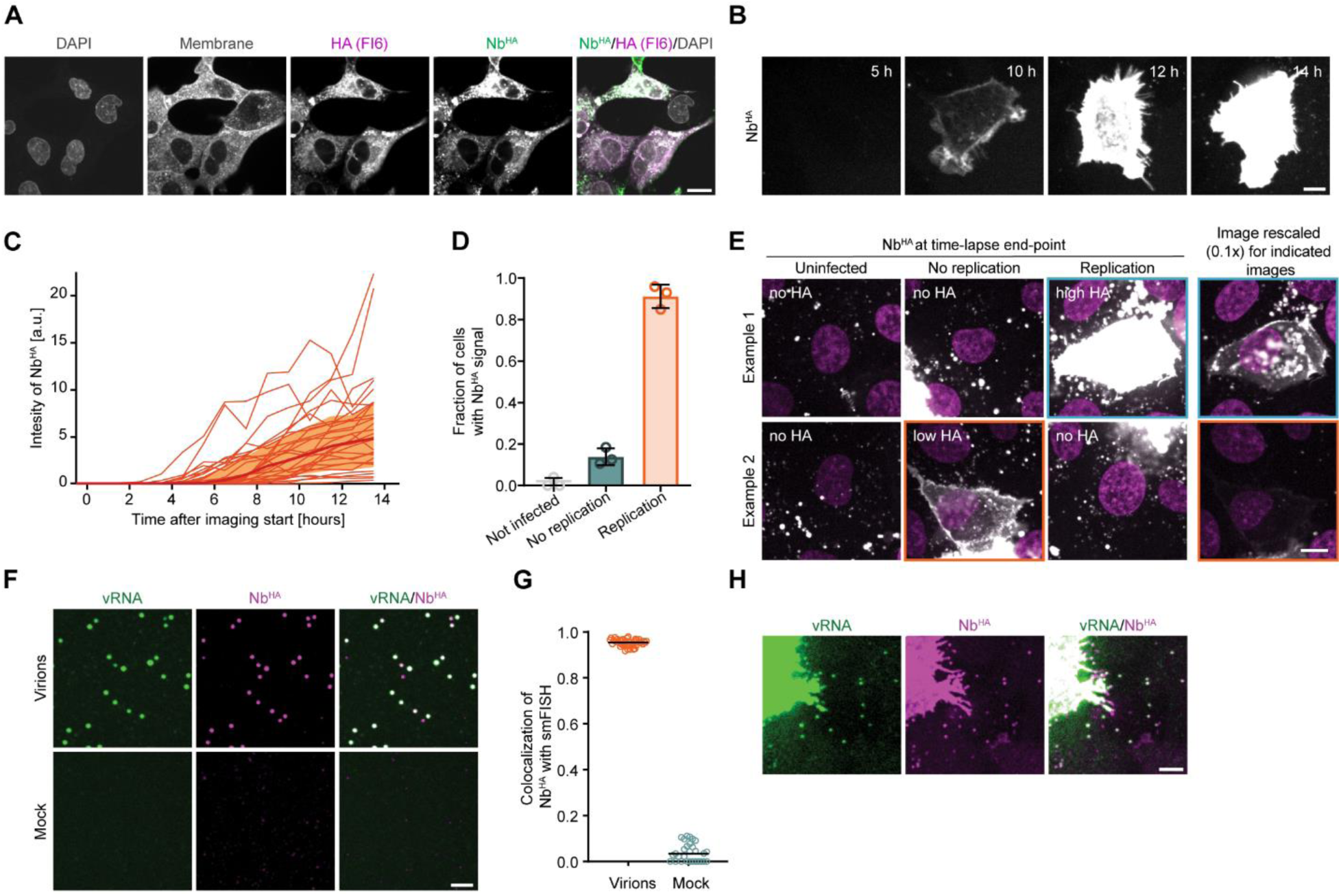
Live-cell extracellular labelling of HA reveals dynamics of viral protein synthesis and progeny production. **(A)** Example images of cells inoculated with IAV for 20 h, fixed and stained with the FI6 anti-HA antibody and the Nb^HA^ nanobody. Scale bar, 15 µm. **(B)** Example images of a time-lapse movie of an IAV infected cell labelled with Nb^HA^. Scale bar, 10 µm. **(C)** Nb^HA^ intensity time traces of single cells that are infected with IAV undergoing replication. Replication was called using VIRIM-neg. **(D)** Quantification of the frequency of Nb^HA^ signal positivity for the indicated infection groups. Dots represent averages of independent experiments. Error bars represent SDs. **(E)** Example images 17.5 hpi of the Nb^HA^ signal in VIRIM-neg cells inoculated with IAV. Two representative cells for each observed phenotype are shown (uninfected, infected with non-replicating virus or infected with replicating virus, as assessed with VIRIM-neg). The orange and the blue image outline indicate the two images that are rescaled in the right column. Scale bar, 10 µm. **(F)** Example images of virions spotted on glass and stained with Nb^HA^ (magenta), as well as smFISH probes targeting all eight vRNAs (green). As control, the supernatant of uninfected cells was used (mock). Scale bar, 5 µm. **(G)** Quantification of colocalization between Nb^HA^ and vRNA smFISH shown in **F**. **(H)** Example images of an infected cell 20 hpi stained with Nb^HA^ and smFISH probes targeting all vRNAs. The outline of the infected cell is visible along with multiple progeny virions (dual-color spots). Scale bar, 5 µm.

**Figure S4.**
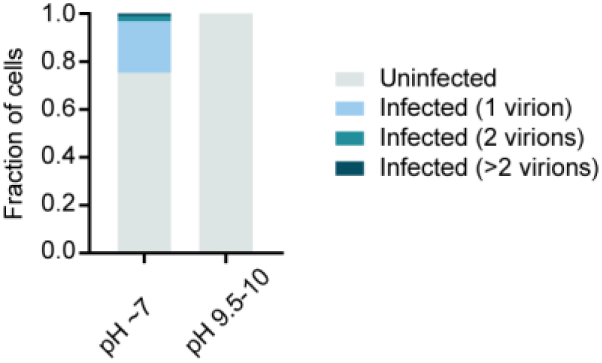
Inhibition of IAV infection in high-pH medium. Quantification of the frequency of infections with 1, 2 or >2 IAV virions, as assessed by VIRIM-neg, in A549-Nb^NP^ cells 1 hpi at 37°C and atmospheric CO_2_. Inoculation was either performed in L15 (pH 7.0-7.4 at atmospheric CO_2_) or in DMEM (pH 9.5-10 at atmospheric CO_2_).

**Figure S5.**
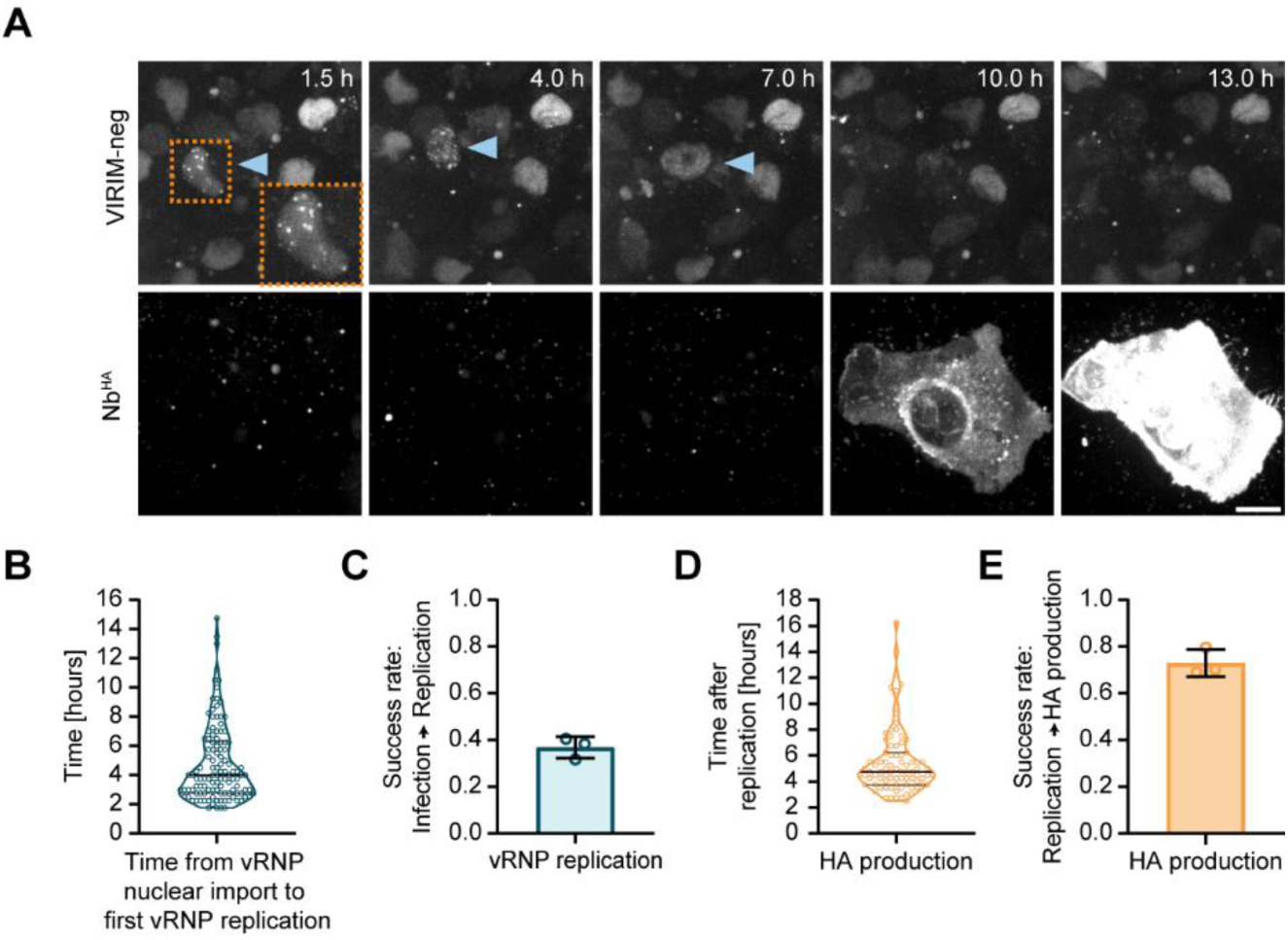
Dissecting the IAV infection cycle in primary airway cells. **(A)** Example images from a time-lapse movie of primary human lung epithelia organoids inoculated with IAV. Images show infection cycle phases as visualized by VIRIM-neg and Nb^HA^. Arrows and zoom-in (orange line) indicate the nucleus of an infected cell. Scale bar, 10 µm. **(B)** Quantification of the time between inoculation with IAV and vRNP replication initiation in lung epithelia organoid cells. **(C)** Quantification of average fraction of infected lung epithelia organoid cells in which vRNP replication was successful. **(D)** Quantification of the time of HA cell surface detection relative to the moment of vRNP replication initiation in lung epithelia organoid cells. **(E)** Quantification of the average fraction of infected lung epithelia organoid cells in which HA protein was detected. **(B,D)** Dots represent individual cells. **(C, E)** Dots represent average values from independent experiments. Error bars represent SDs.

**Figure S6.**
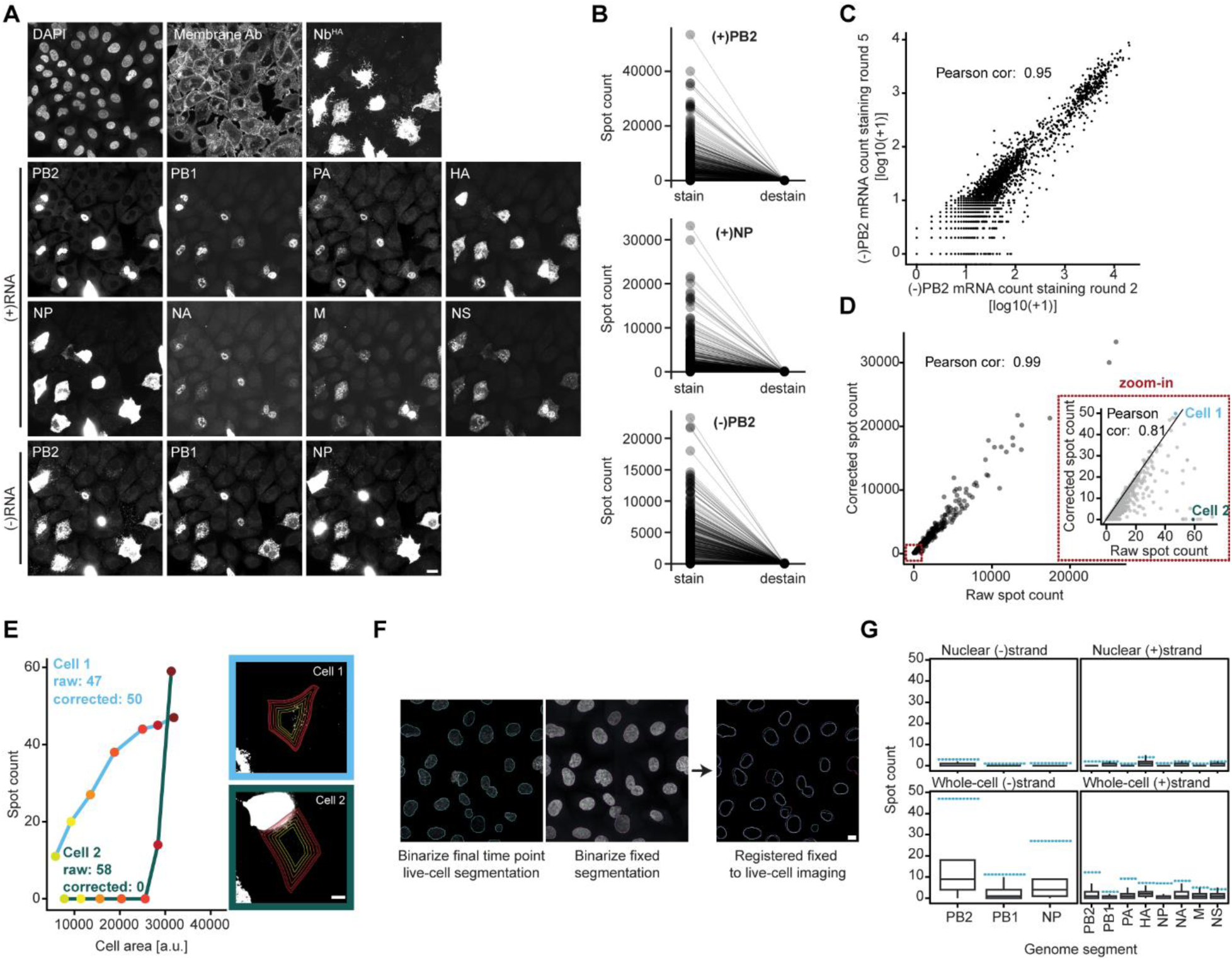
Technical validation of multiplexed smFISH experiments. **(A)** Example images of all smFISH staining rounds, plasma membrane staining, Nb^HA^ and DAPI staining for the same field of view. Scale bar, 20 µm. **(B)** smFISH spot count of (+)PB2, (+)NP or (-)PB2 in single cells before and after destaining. The lines between stain and destain link counts for the same cells. **(C)** Correlation between spot counts of (-)PB2 smFISH detected in staining round 2 and 5. **(D)** Correlation between the raw spot count and the spot count after correction using the iterative erosion approach (see **E** and methods). The red box indicates a zoomed-in area of the graph. Cell 1 is highlighted as an example where spot count is unaffected by spot count correction using the iterative erosion approach, while cell 2 is corrected to zero spots. **(E)** Example images and data for the iterative erosion approach for spot count correction for cell 1 and cell 2 shown in (D). The iterative erosion approach corrects for high smFISH spot counts that are erroneously assigned to cell 2 due to imperfect cell segmentation. Scale bar, 10 µm. **(F)** Example images for the registration process to link live-cell imaging to fixed-cell imaging, Scale bar, 10 µm. **(G)** Spot count in IAV inoculated, but uninfected cells (as determined by VIRIM-neg) is shown as box plots. The 95^th^ percentile of spot counts in uninfected cells for each staining is shown (dashed blue line), and was used as threshold value below which cells were assigned a spot count value of 0.

**Figure S7.**
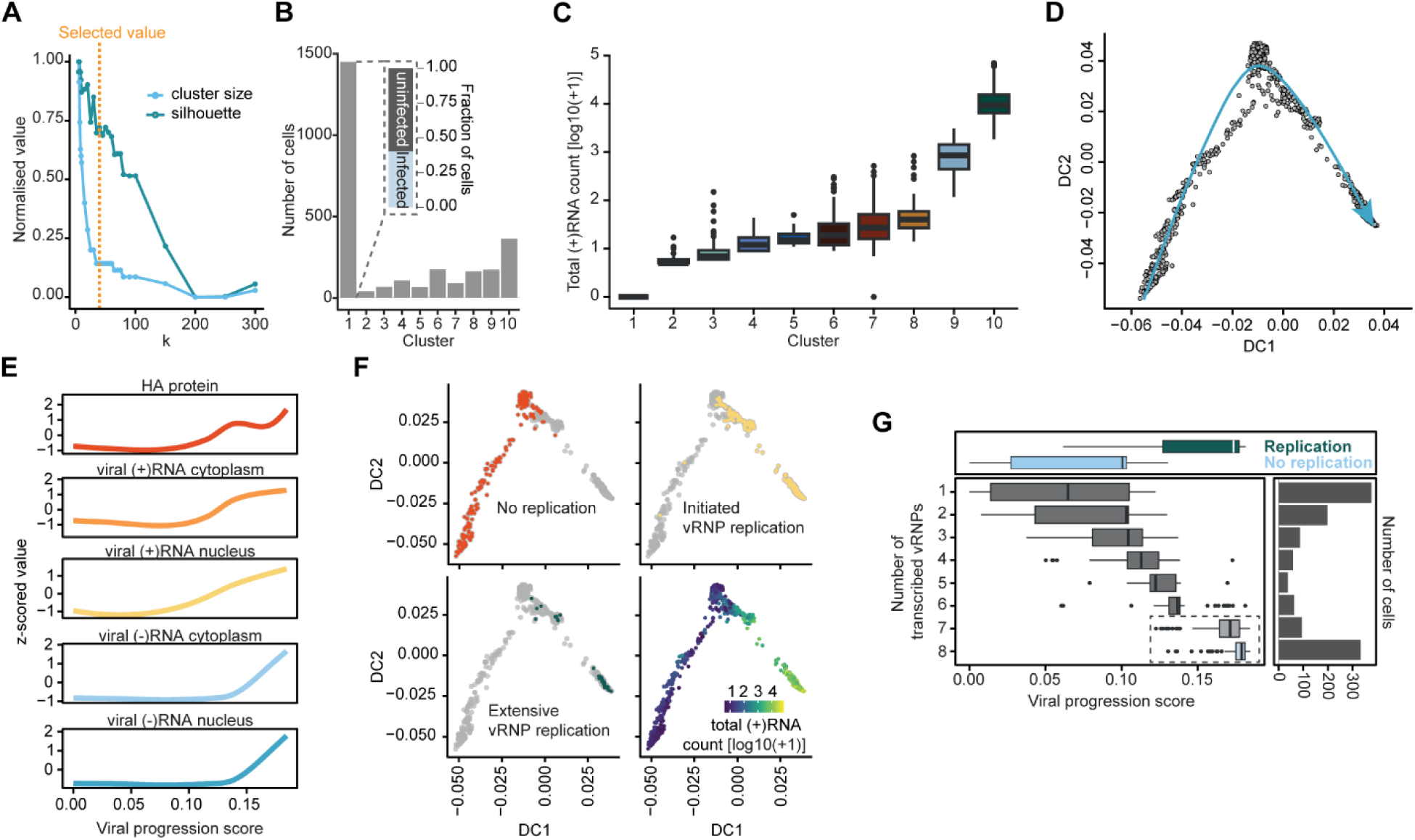
Validation of the multiplexed smFISH approach to assess infection progression. **(A)** Phenograph cluster size and silhouette score as a function of k-parameter in the phenograph algorithm. Selected value (k=40) is indicated with orange dashed line. **(B)** Number of cells in each of the 10 phenograph clusters. Cluster 1 contains cells that lack spots in all smFISH stainings and includes both uninfected cells and infected cells without detectable viral RNA. The fraction of infected and uninfected cells in cluster 1 was estimated by assessing infections in a randomly-selected subset of 278 cluster 1 cells by VIRIM-neg. The estimated fraction of cluster 1 cells that is infected is shown in the zoom-in. **(C)** Sum of all the viral (+)RNA counts in each of the 10 phenograph clusters. **(D)** Diffusion map computed based on: 1) the sum of all eight types of (+)RNA spot counts across nuclei and whole cells, 2) the sum of (-)PB2, (-)PB1, and (-)NP spot counts across nuclei and whole cells and 3) the mean intensity of HA protein across the whole cell. The blue arrow indicates the computed slingshot trajectory along which the progression score for single cells was computed. **(E)** Input features for computing the diffusion map as a function of the final computed viral progression score. **(F)** Diffusion maps as shown in (D). Highlighted are infected cells without replication (top left), infected cells with initiation of replication (top right), or infected cells with extensive replication (bottom left), as determined by VIRIM-neg. The bottom right diffusion map shows the total (+)RNA counts per cell, as determined by smFISH. **(G)** The viral progression score was calculated for each cell (D to F), and cells were grouped by the number of detected viral mRNA species. Viral progression scores for all cells with replicating and non-replicating vRNPs are shown above. Number of cells in each group is shown on the right.

**Figure S8.**
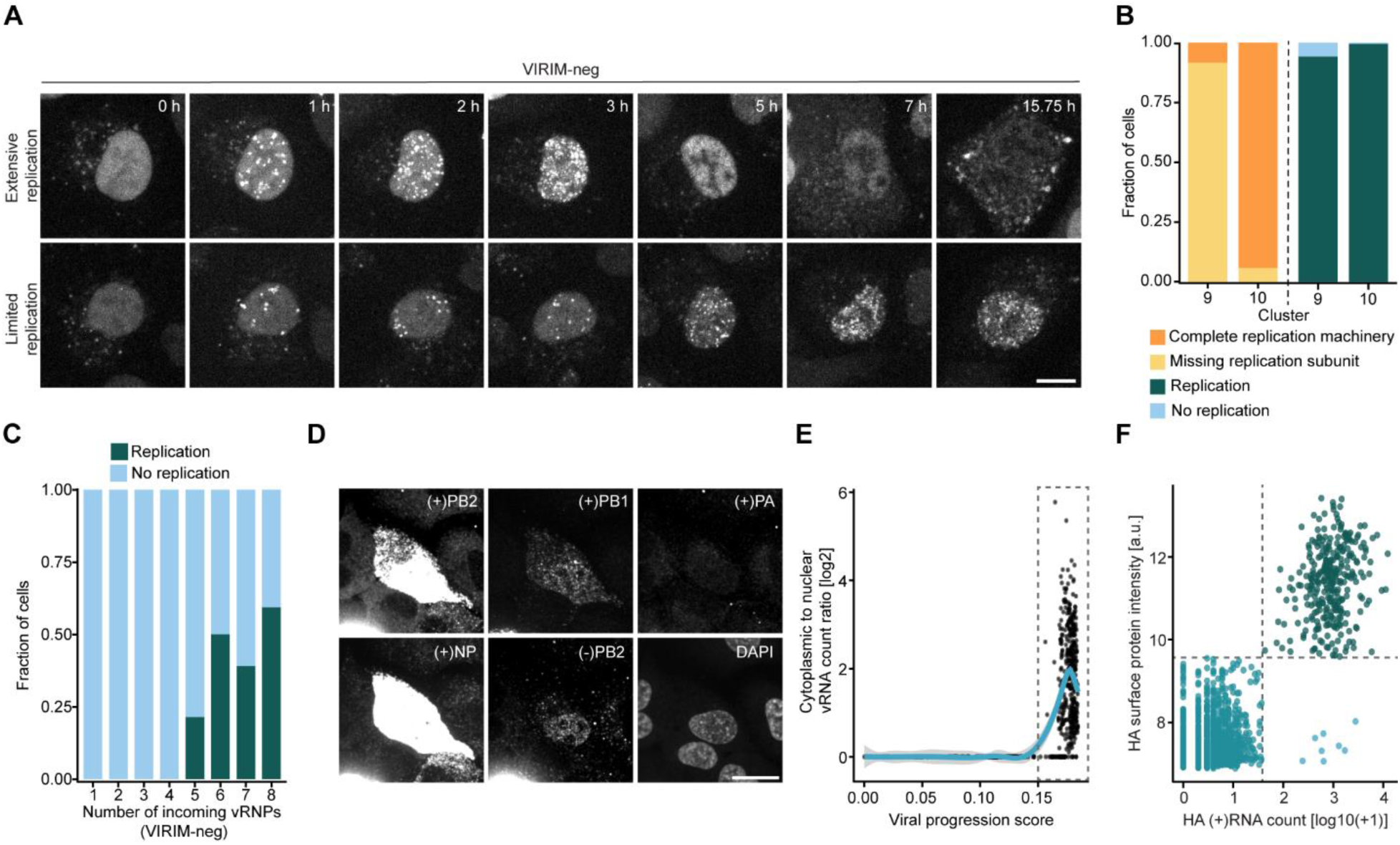
Naturally occurring single-gene knock-out phenotypes in IAV infections. **(A)** Example images of time-lapse movies displaying an infected cell with virus undergoing extensive replication followed by nuclear export of vRNPs (top row), and a cell with virus undergoing limited replication (bottom row). Time indicates hpi. Scale bar, 10 µm. **(B)** Fraction of cells in clusters 9 or 10 that either expressed all four viral proteins essential for replication (PB1, PB2, PA and NP), or that lack expression of one or more of these proteins (left) and fraction of cells in clusters 9 or 10 that showed vRNP replication or not as assessed by VIRIM-neg (right, also displayed in Figure 4I). **(C)** Fraction of cells in which replication occurs. Cells were grouped based on the number of vRNPs delivered into the cell, as determined by VIRIM-neg. **(D)** Ratio between cytoplasmic and nuclear staining intensity of (-)RNAs as a measure of vRNP nuclear export. The box indicates cells for which nuclear export analysis was performed (Figure 4J). **(E)** Correlation of HA surface protein levels, as determined by Nb^HA^ staining, with HA (+)RNA smFISH count in the same cell.

**Figure S9.**
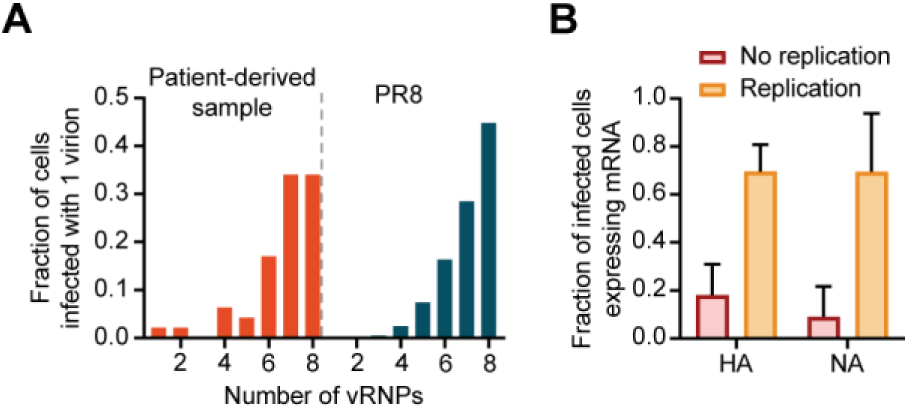
Dissecting the IAV infection cycle of patient-derived virions. **(A)** Quantification of the number of vRNPs per virion for PR8 virus produced in MDCKII cells (blue) and for patient-derived IAV (orange). vRNP count was assessed by VIRIM-neg 1 hpi. Only infections with 8 or less vRNPs were included. **(B)** Quantification of the fraction of patient-derived IAV infected cells expressing HA or NA mRNAs as determined by smFISH. Cells were grouped based on whether genome replication was observed as determined by VIRIM-neg. Error bars represent SDs.

**Figure S10.**
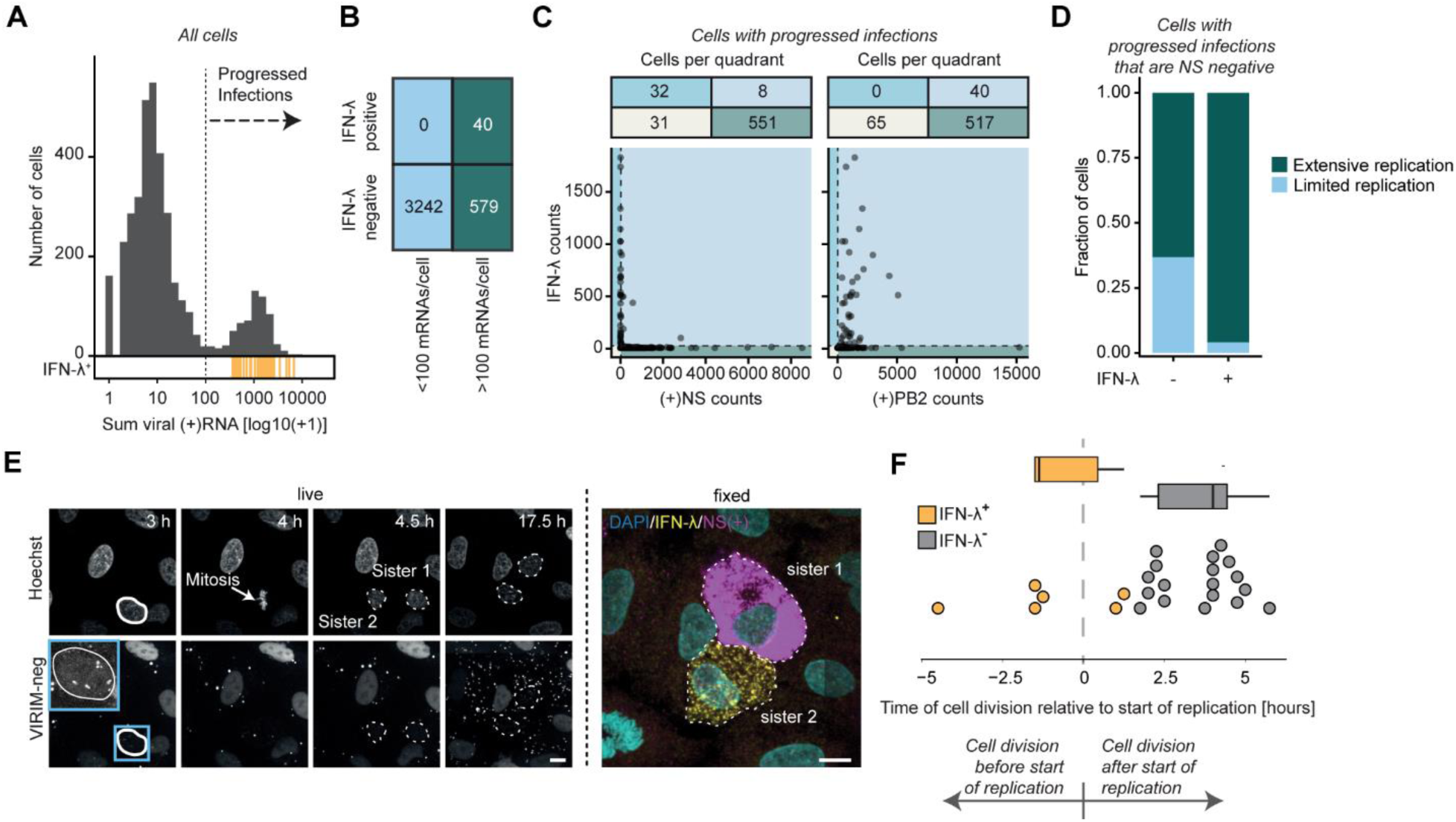
Effects of viral gene expression defects on host immune activation. **(A)** Histogram of the number of cells that contain indicated number of viral mRNAs in the multiplexed smFISH experiment. Infections are considered ‘progressed infections’ if the viral mRNA count exceeds 100 mRNAs per cell (dashed line). Orange lines indicate IFN-λ positive cells. **(B)** Count table showing the number of cells in which IFN-λ mRNA was detected across the groups of uninfected/non-progressed and progressed infections. **(C)** Scatter-plot showing the relationship between IFN-λ and (+)NS or (+)PB2 mRNA counts in cells with progressed infections. Groups were split into quadrants based on the 10^th^ percentile of each count (gray dashed lines). The tables above the graphs indicate the respective cell count in every quadrant (colors in the table refer to the quadrants of the same color). **(D)** Replication status, as assessed by VIRIM-neg, in cells with >100 viral mRNAs (progressed infections) that lack (+)NS mRNA expression. **(E)** Example images of a time-lapse movie of an infected cell that underwent cell division after viral entry. The two sister cells after fixation and smFISH staining for IFN-λ and NS(+) are shown in the right panel. Scale bar, 10 µm. **(F)** IFN-λ expression in relation to the timing of cell division and vRNP replication, as assessed by live-cell imaging. Only infections initiated by a viral particle that contained the NS segment are plotted. Viral particles were called NS-positive if one or both of the sister-cells stained positive for the NS mRNA by smFISH at 19 hpi.

**Link to Videos 1 to 6:**

https://drive.google.com/drive/folders/1k7cNixihQgzYB_msAVsLN3xbtD4b0M6j?usp=sharing

**Video S1.**

**Live-cell imaging of IAV vRNPs by VIRIM-neg**. Imaging of a single GFP z-slice of a A549-Nb^NP^ cell inoculated with IAV 1 h before imaging. Time is indicated as seconds:milliseconds. Video shows an infection by 1 virion that delivered 8 vRNPs into the nucleus. Scale bar, 10 µm.

**Video S2.**

**Time-lapse imaging of extracellular labelling of the HA protein by Nb^HA^ and progeny release from IAV infected cells**. Imaging of 3 z-slices in both the green and blue channels, to visualize Nb^HA^ (shown in white) and cell nuclei (shown in cyan), respectively. Images were acquired every 20 min. Out of 2 cells producing HA protein, only one released virions into the supernatant. Time is indicated as hours:minutes since the start of inoculation. Scale bar is indicated with description throughout the video.

**Video S3.**

**VIRIM-neg imaging of vRNP release into the cytoplasm**. Imaging of a single GFP z-slice of a A549-Nb^NP^ cell 2 min after pH adjustment. Time is indicated as seconds:milliseconds. The cell was infected by 1 virion that delivered 8 vRNPs into the cytoplasm in a step-by-step fashion. Scale bar, 5 µm.

**Video S4.**

**VIRIM-neg imaging of different phenotypes of IAV infected cells**. VIRIM-neg was used to visualize vRNP nuclear entry, vRNP replication and vRNP nuclear export. Images were acquired every 10 min. Images are maximum intensity projections of 3 z-slices. BafA1 was added 1.5 hpi. The first row shows three infections without vRNP replication. The second row shows three examples of infections with vRNP replication but without vRNP nuclear export. The last row shows three examples of cells with successful vRNP replication and nuclear export. Time is indicated as hours:minutes. Scale bar, 25 µm.

**Video S5.**

**VIRIM-neg and Nb^HA^ imaging of different phenotypes of IAV infected cells**. Nb^HA^ was used to visualize cell surface HA protein expression, while VIRIM-neg visualizes vRNPs. Images were acquired every 10 min. Imaging are a maximum intensity projection of 3 z-slices. BafA1 and Nb^HA^ were added 1.5 hpi. Both example cells have vRNP replication. The example cell in the upper row did not show HA protein expression, while the example cell in the bottom row does. Time is indicated as hours:minutes. Scale bar, 25 µm.

**Video S6.**

**Matched VIRIM-neg and Nb^HA^ live-cell imaging with multiplexed smFISH**. Nb^HA^ was used to visualize cell surface HA protein expression, while VIRIM-neg visualizes vRNPs. Initial zoom-in shows three example cells with different infection states: uninfected, infected by a single particle and infected by multiple particles. Zoom-out then shows live-cell time-lapse of viral infection over 19 hpi. Single-cell tracking is visualized by colored tails. 16 hpi cells are fixed and smFISH staining is performed. Multiplexed smFISH data is shown for all eight (+)RNA stainings, first individually and then by merging the stainings in different colors – highlighting the heterogeneity of viral gene expression.

## References

1. Russell, A.B., et al., Single-Cell Virus Sequencing of Influenza Infections That Trigger Innate Immunity. J Virol, 2019. 93(14).

2. Zath, G.K., et al., Influenza A viral burst size from thousands of infected single cells using droplet quantitative PCR (dqPCR). PLoS Pathog, 2024. 20(7): p. e1012257.

3. Russell, A.B., C. Trapnell, and J.D. Bloom, Extreme heterogeneity of influenza virus infection in single cells. Elife, 2018. 7.

4. Kelly, J.N., et al., Comprehensive single cell analysis of pandemic influenza A virus infection in the human airways uncovers cell-type specific host transcriptional signatures relevant for disease progression and pathogenesis. Front Immunol, 2022. 13: p. 978824.

5. Wei, Z., et al., Biophysical characterization of influenza virus subpopulations using field flow fractionation and multiangle light scattering: correlation of particle counts, size distribution and infectivity. J Virol Methods, 2007. 144(1-2): p. 122–32.

6. Donald, H.B. and A. Isaacs, Counts of influenza virus particles. J Gen Microbiol, 1954. 10(3): p. 457–64.

7. Noton, S.L., et al., Studies of an influenza A virus temperature-sensitive mutant identify a late role for NP in the formation of infectious virions. J Virol, 2009. 83(2): p. 562–71.

8. Schmidt, F.I., et al., Phenotypic lentivirus screens to identify functional single domain antibodies. Nat Microbiol, 2016. 1(8): p. 16080.

9. Ochiai, H., et al., Inhibitory effect of bafilomycin A1, a specific inhibitor of vacuolar-type proton pump, on the growth of influenza A and B viruses in MDCK cells. Antiviral Res, 1995. 27(4): p. 425–30.

10. Boersma, S., et al., Translation and Replication Dynamics of Single RNA Viruses. Cell, 2020. 183(7): p. 1930–1945 e23.

11. Davis, A.R., A.L. Hiti, and D.P. Nayak, Influenza defective interfering viral RNA is formed by internal deletion of genomic RNA. Proc Natl Acad Sci U S A, 1980. 77(1): p. 215–9.

12. Gaiotto, T. and S.E. Hufton, Cross-Neutralising Nanobodies Bind to a Conserved Pocket in the Hemagglutinin Stem Region Identified Using Yeast Display and Deep Mutational Scanning. PLoS One, 2016. 11(10): p. e0164296.

13. Caffrey, M. and A. Lavie, pH-Dependent Mechanisms of Influenza Infection Mediated by Hemagglutinin. Front Mol Biosci, 2021. 8: p. 777095.

14. Bonnafous, P. and T. Stegmann, Membrane perturbation and fusion pore formation in influenza hemagglutinin-mediated membrane fusion. A new model for fusion. J Biol Chem, 2000. 275(9): p. 6160–6.

15. Qin, C., et al., Real-time dissection of dynamic uncoating of individual influenza viruses. Proc Natl Acad Sci U S A, 2019. 116(7): p. 2577–2582.

16. Chou, Y.Y., et al., Colocalization of different influenza viral RNA segments in the cytoplasm before viral budding as shown by single-molecule sensitivity FISH analysis. PLoS Pathog, 2013. 9(5): p. e1003358.

17. Heldt, F.S., et al., Single-cell analysis and stochastic modelling unveil large cell-to-cell variability in influenza A virus infection. Nat Commun, 2015. 6: p. 8938.

18. Sachs, N., et al., Long-term expanding human airway organoids for disease modeling. EMBO J, 2019. 38(4).

19. York, A., et al., Isolation and characterization of the positive-sense replicative intermediate of a negative-strand RNA virus. Proc Natl Acad Sci U S A, 2013. 110(45): p. E4238–45.

20. Fan, H., et al., Structures of influenza A virus RNA polymerase offer insight into viral genome replication. Nature, 2019. 573(7773): p. 287-290.

21. Brunotte, L., et al., The nuclear export protein of H5N1 influenza A viruses recruits Matrix 1 (M1) protein to the viral ribonucleoprotein to mediate nuclear export. J Biol Chem, 2014. 289(29): p. 20067–77.

22. Bui, M., et al., Role of the influenza virus M1 protein in nuclear export of viral ribonucleoproteins. J Virol, 2000. 74(4): p. 1781–6.

23. O’Neill, R.E., J. Talon, and P. Palese, The influenza virus NEP (NS2 protein) mediates the nuclear export of viral ribonucleoproteins. EMBO J, 1998. 17(1): p. 288–96.

24. Martin, K. and A. Helenius, Nuclear transport of influenza virus ribonucleoproteins: the viral matrix protein (M1) promotes export and inhibits import. Cell, 1991. 67(1): p. 117–30.

25. Marjuki, H., et al., Membrane Accumulation of Influenza A Virus Hemagglutinin Triggers Nuclear Export of the Viral Genome via Protein Kinase Cα-mediated Activation of ERK Signaling. Journal of Biological Chemistry, 2006. 281(24): p. 16707–16715.

26. Nakatsu, S., et al., Complete and Incomplete Genome Packaging of Influenza A and B Viruses. mBio, 2016. 7(5).

27. Chou, Y.Y., et al., One influenza virus particle packages eight unique viral RNAs as shown by FISH analysis. Proc Natl Acad Sci U S A, 2012. 109(23): p. 9101–6.

28. Schelker, M., et al., Viral RNA Degradation and Diffusion Act as a Bottleneck for the Influenza A Virus Infection Efficiency. PLoS Comput Biol, 2016. 12(10): p. e1005075.

29. Le Sage, V., et al., Mapping of Influenza Virus RNA-RNA Interactions Reveals a Flexible Network. Cell Rep, 2020. 31(13): p. 107823.

30. Noda, T., et al., Three-dimensional analysis of ribonucleoprotein complexes in influenza A virus. Nat Commun, 2012. 3: p. 639.

31. Sun, J., et al., Single cell heterogeneity in influenza A virus gene expression shapes the innate antiviral response to infection. PLoS Pathog, 2020. 16(7): p. e1008671.

32. Killip, M.J., et al., Single-cell studies of IFN-beta promoter activation by wild-type and NS1-defective influenza A viruses. J Gen Virol, 2017. 98(3): p. 357–363.

33. Vicary, A.C., et al., Maximal interferon induction by influenza lacking NS1 is infrequent owing to requirements for replication and export. PLoS Pathog, 2023. 19(4): p. e1010943.

34. Ramos, I., et al., Innate Immune Response to Influenza Virus at Single-Cell Resolution in Human Epithelial Cells Revealed Paracrine Induction of Interferon Lambda 1. J Virol, 2019. 93(20).

35. Liu, Y.G., et al., Interferon lambda in respiratory viral infection: immunomodulatory functions and antiviral effects in epithelium. Front Immunol, 2024. 15: p. 1338096.

36. Yang, H., et al., The influenza virus PB2 protein evades antiviral innate immunity by inhibiting JAK1/STAT signalling. Nat Commun, 2022. 13(1): p. 6288.

37. Graef, K.M., et al., The PB2 subunit of the influenza virus RNA polymerase affects virulence by interacting with the mitochondrial antiviral signaling protein and inhibiting expression of beta interferon. J Virol, 2010. 84(17): p. 8433–45.

38. Iwai, A., et al., Influenza A virus polymerase inhibits type I interferon induction by binding to interferon beta promoter stimulator 1. J Biol Chem, 2010. 285(42): p. 32064–74.

39. de Wit, E., et al., Efficient generation and growth of influenza virus A/PR/8/34 from eight cDNA fragments. Virus Res, 2004. 103(1-2): p. 155–61.

40. Lambert, G.G., et al., Aequorea’s secrets revealed: New fluorescent proteins with unique properties for bioimaging and biosensing. PLoS Biol, 2020. 18(11): p. e3000936.

41. Ivorra-Molla, E., et al., A monomeric StayGold fluorescent protein. Nat Biotechnol, 2024. 42(9): p. 1368–1371.

42. Hufton, S.E., et al., The breadth of cross sub-type neutralisation activity of a single domain antibody to influenza hemagglutinin can be increased by antibody valency. PLoS One, 2014. 9(8): p. e103294.

43. Zeng, Q., et al., Structure of coronavirus hemagglutinin-esterase offers insight into corona and influenza virus evolution. Proc Natl Acad Sci U S A, 2008. 105(26): p. 9065–9.

44. Dayton, T.L., et al., Druggable growth dependencies and tumor evolution analysis in patient-derived organoids of neuroendocrine neoplasms from multiple body sites. Cancer Cell, 2023. 41(12): p. 2083–2099 e9.

45. Liu, M.Y., et al., Human Airway and Alveolar Organoids from BAL Fluid. Am J Respir Crit Care Med, 2024. 209(12): p. 1501–1504.

46. Wijnakker, J., et al., Integrin-activating Yersinia protein Invasin sustains long-term expansion of primary epithelial cells as 2D organoid sheets. Proc Natl Acad Sci U S A, 2025. 122(1): p. e2420595121.

47. Lei, C., et al., On the Calculation of TCID(50) for Quantitation of Virus Infectivity. Virol Sin, 2021. 36(1): p. 141–144.

48. Fouchier, R.A., et al., Detection of influenza A viruses from different species by PCR amplification of conserved sequences in the matrix gene. J Clin Microbiol, 2000. 38(11): p. 4096–101.

49. Gaspar, I., F. Wippich, and A. Ephrussi, Terminal Deoxynucleotidyl Transferase Mediated Production of Labeled Probes for Single-molecule FISH or RNA Capture. Bio Protoc, 2018. 8(5): p. e2750.

50. Lyubimova, A., et al., Single-molecule mRNA detection and counting in mammalian tissue. Nat Protoc, 2013. 8(9): p. 1743–58.

51. Corti, D., et al., A neutralizing antibody selected from plasma cells that binds to group 1 and group 2 influenza A hemagglutinins. Science, 2011. 333(6044): p. 850-6.

52. Chalfoun, J., et al., MIST: Accurate and Scalable Microscopy Image Stitching Tool with Stage Modeling and Error Minimization. Sci Rep, 2017. 7(1): p. 4988.

53. Schindelin, J., et al., Fiji: an open-source platform for biological-image analysis. Nat Methods, 2012. 9(7): p. 676-82.

54. Stringer, C., et al., Cellpose: a generalist algorithm for cellular segmentation. Nat Methods, 2021. 18(1): p. 100–106.

55. Ershov, D., et al., TrackMate 7: integrating state-of-the-art segmentation algorithms into tracking pipelines. Nat Methods, 2022. 19(7): p. 829–832.

56. Eichenberger, B.T., et al., deepBlink: threshold-independent detection and localization of diffraction-limited spots. Nucleic Acids Res, 2021. 49(13): p. 7292–7297.

57. Peng, T., et al., A BaSiC tool for background and shading correction of optical microscopy images. Nat Commun, 2017. 8: p. 14836.

58. Thevenaz, P., U.E. Ruttimann, and M. Unser, A pyramid approach to subpixel registration based on intensity. IEEE Trans Image Process, 1998. 7(1): p. 27–41.

59. Pachitariu, M. and C. Stringer, Cellpose 2.0: how to train your own model. Nat Methods, 2022. 19(12): p. 1634–1641.

60. Sofroniew, N., Lambert, T., Bokota, G., Nunez-Iglesias, J., Sobolewski, P., Sweet, A., Gaifas, L., Evans, K., Burt, A., Doncila Pop, D., Yamauchi, K., Weber Mendonça, M., Buckley, G., Vierdag, W.-M., Royer, L., Can Solak, A., Harrington, K. I. S., Ahlers, J., Althviz Moré, D., …, Zhao, R., napari: a multi-dimensional image viewer for Python (v0.5.4). 2024, Zenodo.

61. Imbert, A., et al., FISH-quant v2: a scalable and modular tool for smFISH image analysis. RNA, 2022. 28(6): p. 786–795.

62. van der Walt S, S.J., Nunez-Iglesias J, Boulogne F, Warner JD, Yager N, Gouillart E, Yu T, the scikit-image contributors, scikit-image: image processing in Python. PeerJ, 2014. 2:e453.

63. Team, R.C., A language and environment for statistical computing.. 2022, R Foundation for Statistical Computing: Vienna, Austria.

64. Wickham, H., ggplot2: Elegant Graphics for Data Analysis. 2016: Springer-Verlag New York.

65. Wickham H, F.R., Henry L, Müller K, Vaughan D, dplyr: A Grammar of Data Manipulation. 2023.

66. Levine, J.H., et al., Data-Driven Phenotypic Dissection of AML Reveals Progenitor-like Cells that Correlate with Prognosis. Cell, 2015. 162(1): p. 184–97.

67. McInnes, L.H., J.; Melville, J., UMAP: Uniform Manifold Approximation and Projection for Dimension Reduction. arXiv, 2020.

68. Angerer, P., et al., destiny: diffusion maps for large-scale single-cell data in R. Bioinformatics, 2016. 32(8): p. 1241–3.

69. Street, K., et al., Slingshot: cell lineage and pseudotime inference for single-cell transcriptomics. BMC Genomics, 2018. 19(1): p. 477.

